# The role of water mobility in protein misfolding

**DOI:** 10.1101/2021.01.06.425575

**Authors:** Amberley D. Stephens, Johanna Kölbel, Rani Moons, Michael T. Ruggerio, Najet Mahmoudi, Talia A. Shmool, Thomas M. McCoy, Daniel Nietlispach, Alexander F. Routh, Frank Sobott, J. Axel Zeitler, Gabriele S. Kaminski Schierle

## Abstract

The propensity for intrinsically disordered proteins to aggregate is heavily influenced by their surrounding environment. Here, we show that the mobility of the surrounding water molecules directly influences the aggregation rate of α-synuclein (aSyn), a protein associated with Parkinson’s disease. We observe that the addition of NaCl reduces the mobility of water, while addition of CsI increases the mobility of water. In turn, this reduces and increases the mobility of aSyn, respectively, given the change in strength and lifetime of the intermolecular forces. The reduction of aSyn mobility in the presence of NaCl ions leads to increased aggregation rates, which may be due to aggregation-competent conformations being stable for longer, thereby increasing the likelihood of establishing interactions between two adjacent monomers. In contrast, aSyn is more mobile when CsI is dissolved in the aqueous phase which leads to a reduction of successful monomeric interactions. We thus highlight the importance of the surrounding environment and describe how ion content can influence water mobility and the misfolding rate of amyloidogenic proteins, such as aSyn. By modulating the cellular environment to increase water mobility or finding small molecules to increase protein dynamics, new therapeutic targets may be found.

## Introduction

The majority of proteins cannot function without a solvation shell, and the mobility of this solvation layer affects rates of conformational change, catalysis and protein/DNA-protein interactions^1–4^. Solvent interaction is particularly pertinent for intrinsically disordered proteins (IDPs) which have large solvent accessible areas compared to globular proteins of a similar size^5^. However, it is not currently clear what role the solvent plays in the misfolding and aggregation of proteins, particularly for IDPs such as α-synuclein (aSyn), whose aggregation is a hallmark of synucleinopathies such as Parkinson’s disease. Certainly, water molecules are expelled from the solvation shell for monomer-monomer interactions, fibril elongation and fibril bundling to occur^6^. Water is an important driving force for protein folding, but a detailed understanding of the role of water in stabilising IDPs or in destabilising the protein to influence misfolding and its aggregation propensity is yet to be achieved. Furthermore, it is well-known that ions influence the hydrogen bond dynamics of water molecules^7–12^. Despite this being well-known, the influence of salt ions on water mobility within differing cellular environments, and the subsequent impact this can have on protein misfolding, is currently not fully understood. Here, we show that the addition of NaCl, comprising two small, high charge density ions, and CsI, comprising two large, low charge density ions, can increase and decrease the aggregation rate of aSyn, respectively. Water and aSyn mobility are inextricably linked and increasing water mobility with addition of CsI subsequently increases the protein mobility which reduces the propensity of aSyn to aggregate.

## Results

### CsI decreases aSyn aggregation rate whereas NaCl and D_2_O increase aSyn aggregation rate

Fibrillisation rates of aSyn in the presence of NaCl and CsI were monitored using a fluorescence based kinetic assay with the molecule Thioflavin-T (ThT) which fluoresces when intercalated into the backbone of β-sheet containing fibrils^13,14^. The sigmoidal kinetic curves, representative of nucleation dependent reactions, show that aggregation of aSyn occurred faster in the presence of NaCl compared to CsI in H_2_O (Figure 1a, Supplementary Figure 1). Furthermore, upon increasing concentration of NaCl from 150 mM to 1.5 M, the aSyn aggregation rate increased, as lag time (*t*_lag_) and time to reach half maximum fluorescence (*t*_50_) both decreased (Table 1). Conversely, aggregation of aSyn in the presence of CsI was slower at 1.5 M than at 150 mM concentrations and significantly slower compared to NaCl.

**Figure 1.**
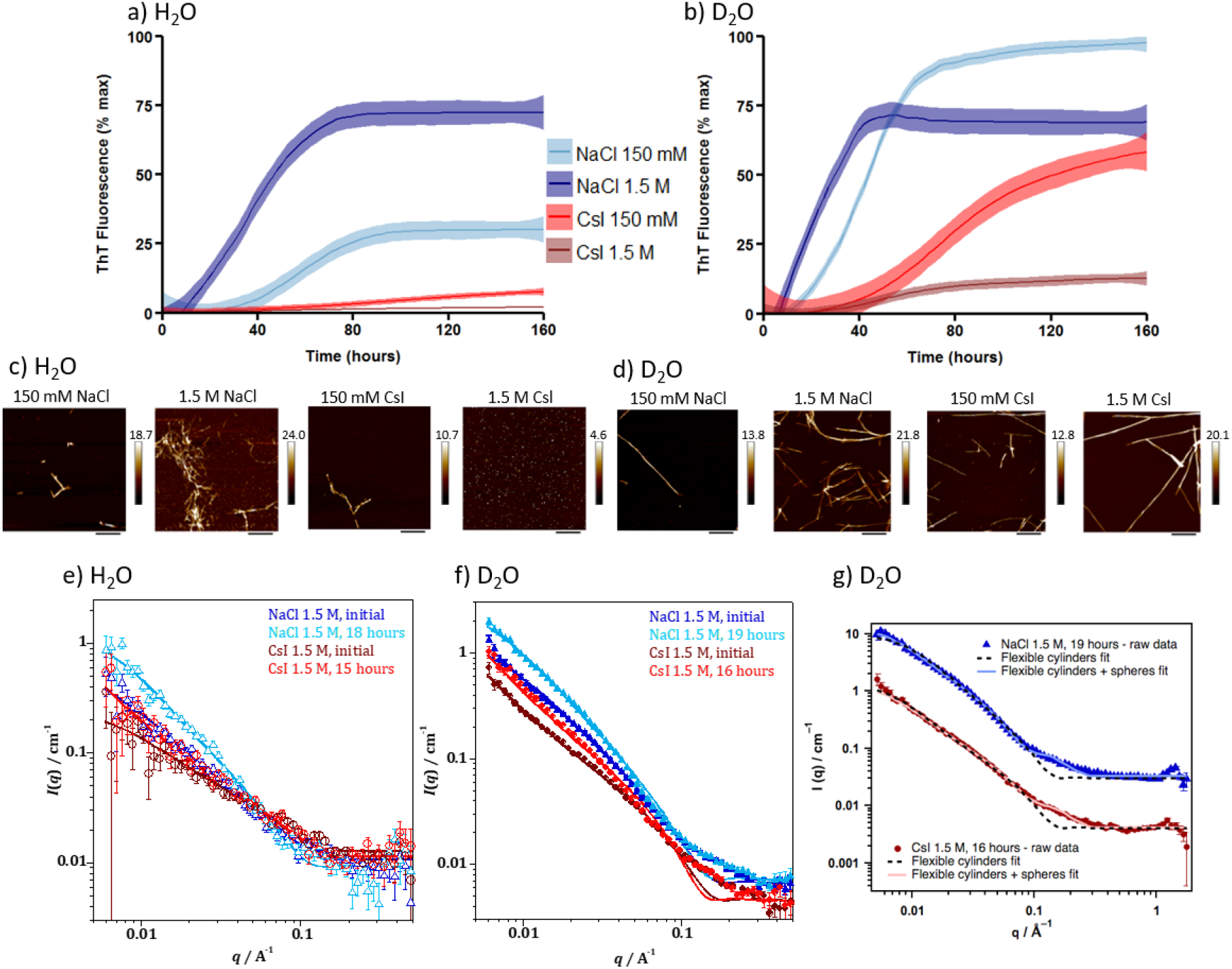
NaCl and CsI concentrations influence aSyn aggregation rate and morphology. aSyn aggregation kinetics were measured in the presence of (a) H_2_O and (b) D_2_O with 150 mM NaCl (red), 1.5 M NaCl (brown), 150 mM CsI (light blue), 1.5 M CsI (navy) and plotted as % maximum ThT fluorescence over time (Supplementary Figure 1 displays individual plate repeats). Data represent three experiments with three or four wells per condition per experiment; error (shaded areas) represents rolling average of the SEM. After the ThT-based assays, aSyn was incubated on freshly cleaved mica and representative images are shown for aSyn species formed in the presence of NaCl and CsI at 150 mM and 1.5 M in (c) H_2_O and (d) D_2_O. Scale bar = 800 nm. For SANS measurements a high concentration (434 μM) of aSyn was used to ensure a sufficient number of scatter counts were attained. Model fits to the SANS data, using a flexible cylinder model, of aSyn in 1.5 M CsI and NaCl in (e) H_2_O and (f) D_2_O after initial mixing and incubation for 15-19 hours. (g) Fittings to a flexible cylinder with spheres (pale filled line) described more accurately the data than fitting to a flexible cylinders model only (dashed line) using data from aSyn in 1.5 M salts in D_2_O. The NaCl (light blue) is offset by a factor of 10 for clarity.

**Table 1.** Lag time (*t*_lag_), time to reach half maximum fluorescence (*t*_50_) and remaining monomer concentration determined by SEC after performing ThT-based kinetic assays

| Solvent | Salt concentration | $t_{lag}$ (hours) | $t_{50}$ (hours) | Remaining Monomer (μM) | % max fluorescence at 160 hours |
| --- | --- | --- | --- | --- | --- |
| H<sub>2</sub>O | NaCl 150 mM | 34.3 ± 4.9 | 47.3 ± 11.5 | 12.8 ± 9.0 | 30.3 ± 14.9 |
| | NaCl 1.5 M | 31.0 ± 5.1 | 41.0 ± 7.9 | 0 | 72.6 ± 11.5 |
| | CsI 150 mM | nd | nd | 38.1 ± 7.8 | 7.9 ± 6.5 |
| | CsI 1.5 M | nd | nd | 35.9 ± 16.6 | 2.1 ± 1.0 |
| D<sub>2</sub>O | NaCl 150 mM | 23.8 ± 4.0 | 45.3 ± 5.2 | 2.0 ± 3.0 | 98.1 ± 0.9 |
| | NaCl 1.5 M | 19.0 ± 2.6 | 34.7 ± 7.2 | 0 | 69.0 ± 15.3 |
| | CsI 150 mM | 39.3 ± 0.9 | 86.3 ± 6.3 | 11.7 ± 2.7 | 59.1 ± 25.1 |
| | CsI 1.5 M | nd | nd | 0 | 12.8 ± 8.2 |

<sup>nd</sup> not determined due to lack of clear ThT response to allow calculations.

In order to further probe the influence of the solvent on aSyn aggregation, the same experiment was performed in a D_2_O solvated environment. We observed the same trends as for the H_2_O samples, i.e. the aggregation rate increased upon addition of NaCl, but decreased upon addition of CsI. Yet, in all samples, the substitution of H_2_O for D_2_O accelerated the aSyn aggregation rate (Figure 1a,b, Table 1). We investigated the morphology of the resulting aSyn aggregates (Figure 1c,d, Supplementary Figure 2) and the extent of aggregation by the quantity of remaining monomer after the kinetic assays (Table 1, Supplementary Figure 3). The results mostly reflected the observed aggregation endpoints of the ThT-based assays, but in the CsI containing samples oligomeric species were detected using atomic force microscopy (AFM) and size exclusion chromatography (SEC) (Figure 1c, Supplementary Figure 3), and these species may not bind ThT^15^.

To investigate early time points in the aggregation pathway, when formation of oligomers cannot be detected using ThT fluorescence, we used small angle neutron scattering (SANS) to evaluate size and structure differences of aSyn species. SANS data show that, even at early time points, ^~^15-19 hours, aSyn species of larger sizes are already present in NaCl containing solutions (Figure 1e-g, Supplementary Tables 1 and 2). Using different fitting parameters to analyse the SANS data (discussed in Supplementary Note 1) we show that there is four times more monomeric aSyn, classified as a sphere and compared to cylindrical fibrillar structures, present in 1.5 M CsI samples compared to three times more monomeric aSyn in 1.5 M NaCl in D_2_O samples (Table 2, Supplementary Figure 4). The combined results suggest that CsI reduces the aggregation rate of aSyn compared to NaCl and that D_2_O increases the aggregation rate of aSyn compared to H_2_O.

**Table 2.**
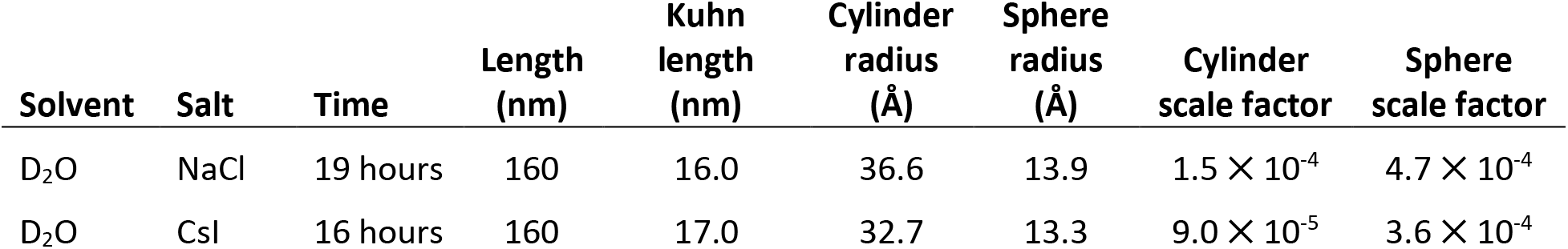
Parameters of fitting SANS data presented in Figures 2g. These results were obtained using a flexible cylinder and sphere model where sphere represents monomeric structures and cylinder fibrillar structures

| Solvent | Salt | Time | Length (nm) | Kuhn length (nm) | Cylinder radius (Å) | Sphere radius (Å) | Cylinder scale factor | Sphere scale factor |
| --- | --- | --- | --- | --- | --- | --- | --- | --- |
| D<sub>2</sub>O | NaCl | 19 hours | 160 | 16.0 | 36.6 | 13.9 | $1.5 \times 10^{-4}$ | $4.7 \times 10^{-4}$ |
| D<sub>2</sub>O | CsI | 16 hours | 160 | 17.0 | 32.7 | 13.3 | $9.0 \times 10^{-5}$ | $3.6 \times 10^{-4}$ |

### MD simulations show solvated water and aSyn_72-78_ peptide mobility is increased in the presence of CsI, but reduced in the presence of NaCl

Ab initio molecular dynamics (AIMD) simulations can be used to elucidate the dynamics of the solvation shell(s), ions, as well as of aSyn^16^ on fs to ps timescales. The simulation results performed on a crystal structure of seven amino acids (TGVGAGA, residues 72-78) from the central region of aSyn highlight that the average velocity of the respective particles remain relatively constant across the various simulations. However, the diffusion constants show that the CsI system results in a significantly increased diffusion (i.e. further displacement from initial positions) of all of the components of the system, and results in a diffusion constant of the aSyn_72-78_ peptide of more than double that determined for the pure water and NaCl models (Table 3, Figure 2a, Supplementary Video 1 and 2). This occurs through significant disruption of the water molecules near the coordination sphere of the Cs^+^ cation. Due to its large size, a significant perturbation to the water geometries was observed. As a result, large-scale reorganisation of the water molecules occurs upon motion of Cs^+^ ions which in turn leads to considerable shuttling of the water molecules. Given the strong intermolecular interactions as a result of dispersion, dipole-dipole, hydrogen bonding, and ion-dipole interactions between the water and aSyn_72-78_ peptide molecules this perturbation is coupled strongly to the aSyn_72-78_ peptide molecule and affects the aSyn_72-78_ peptide motions to a large spatial extent. It is important to note that in both the NaCl and CsI simulations, the number of ions within the first solvation shell of the aSyn_72-78_ peptide remain consistent. These data indicate that the presence of these ions could have a great effect on both the water mobility in the solvation shell and protein mobility *in vitro*. The rate of dimerisation is directly linked to the rate of protein reconfiguration, where slow reconfiguration allows for dimerisation to occur, while fast reconfiguration reduces the likelihood of sustainable contacts that result in successful dimerisation^17^.

**Table 3.** Diffusion coefficient and average velocity of the aSyn_72-78_ peptide, solvated water and non-solvated water calculated using MD simulations

| | | Diffusion Coefficient<br>($\times 10^{-9} \text{ m}^2/\text{s}$) | Average Velocity<br>(m/s) |
| --- | --- | --- | --- |
| aSyn<sub>72-78</sub> peptide | Water | 193.8 | 1595.8 |
|  | NaCl | 195.3 | 1671.9 |
|  | CsI | 402.6 | 1609.5 |
| Solvated Water | Water | 550.3 | 1805.6 |
|  | NaCl | 475.5 | 1836.6 |
|  | CsI | 860.4 | 1843.3 |
| Non-Solvated Water | Water | 655.5 | 1740.2 |
|  | NaCl | 667.4 | 1792.5 |
|  | CsI | 811.2 | 1767.4 |

**Figure 2.**
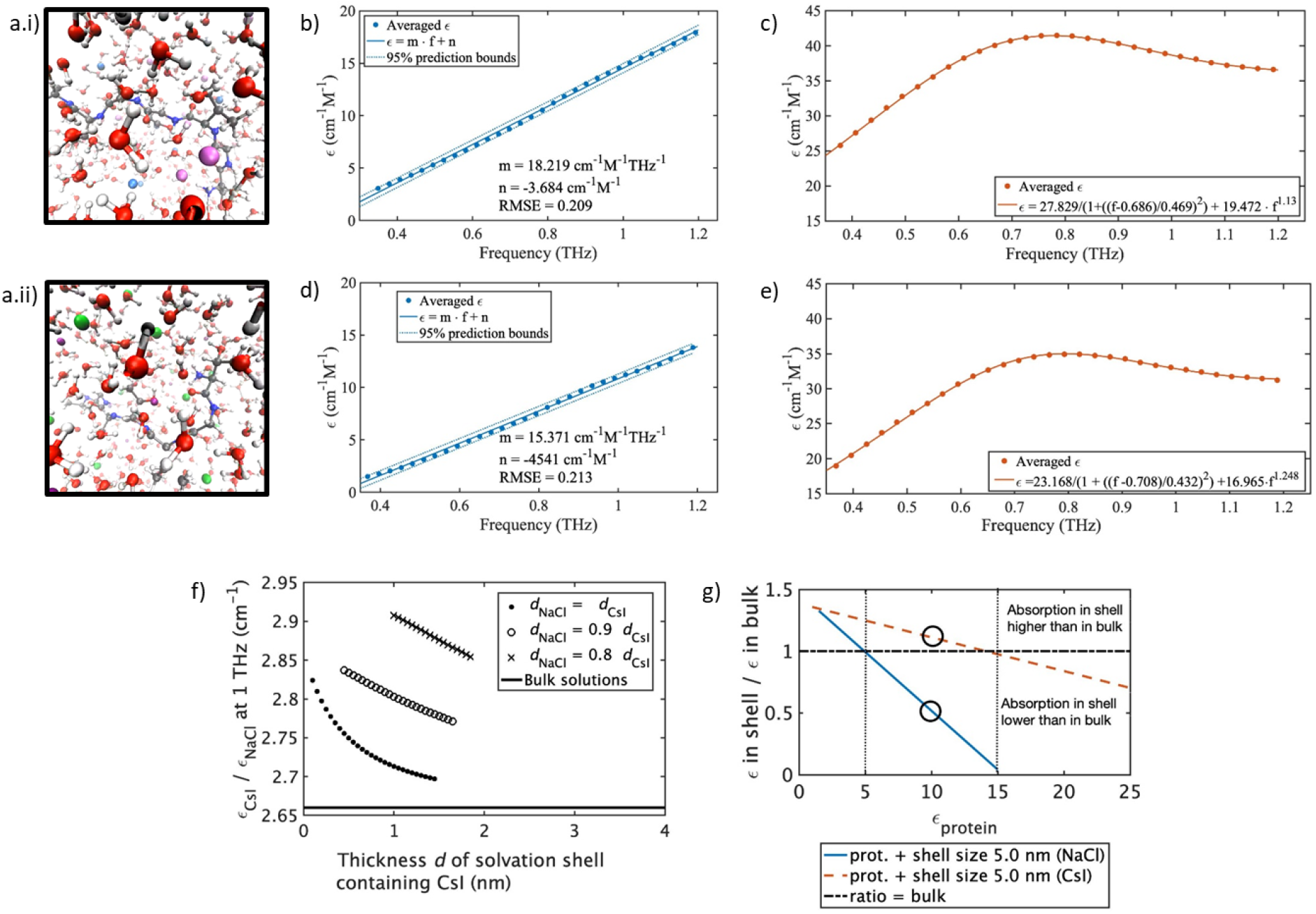
Addition of NaCl and CsI alter water mobility in the bulk and in the aSyn solvation shell. A snapshot of the AIMD simulations of the solvated aSyn_72-78_ peptide in a 125 nm^3^ box after introduction and equilibration with 1.5 M salts. (a.i) aSyn_72-78_ peptide in 1.5 M CsI and (a.ii) aSyn_72-78_ peptide in 1.5 M NaCl, Cs^+^ light purple, I^-^ light blue, Na^+^ dark purple, Cl^-^ green, O red, H white, C grey, N dark blue. The molar absorption coefficient measured with THz-TDS for (b) NaCl solutions, (c) CsI solutions, (d) solutions of NaCl and aSyn, and (e) solutions of CsI and aSyn. (b) and (d) are fitted with a linear function and (c) and (e) with the sum of a power law and a Laurentzian to account for the spectral shape. (f) At 1 THz the solvation shell surrounding aSyn containing CsI (*ε*_CsI_) absorbs more than the shell containing NaCl (ε_NaCl_) and both solvation shells absorb more compared to bulk only (black line at 2.66). The dependency of absorption on the salt is plotted for several possible sizes *d* of the solvation shell, allowing a smaller shell size *d*_NaCl_ in solutions containing NaCl. The same trend is apparent for all shell sizes. (g) Absorption in the solvation shell excluding the protein itself is compared to bulk absorption at 1 THz. Representative ratio of the absorption in the shell compared to bulk absorption for NaCl (blue line) and CsI (orange, dotted) for varying protein absorption (*ε*_P_) between 1-25 cm^−1^M^−1^. The solvation shell with NaCl absorbs less than the bulk (denoted by the black dashed horizontal line) above *ε*_P_ =5 cm^−1^M^−1^ (denoted by the black vertical dotted line at 5 cm^−1^M^−1^), while the solvation shell with CsI absorbs less than the bulk above *ε*_P_ = 15 cm^−1^M^−1^ (denoted by the black dotted vertical line 15 cm^−1^M^−1^) at an ion shell size of 5 nm. At *ε*_P_ = 10 cm^−1^M^−1^, an intermediate protein absorption coefficient, the shell absorption in NaCl is reduced while increased in CsI (shown by black circles). Other shell sizes are shown in Supplementary Figure 6.

### The mobility of water increases in bulk and in the solvation shell in the presence of CsI compared to NaCl

Terahertz time-domain spectroscopy can be used as a highly sensitive probe for water mobility in the liquid state^18,19^. Using THz-TDS measurements in the absence of protein we observe a larger overall increase in absorption coefficient for solutions containing CsI than in the ones containing NaCl (Figure 2b,c). In line with previous results, samples with protein lead to a reduced absorption coefficient as the protein displaces the ions and water molecules which have much stronger absorption than the protein due to the relative number of oscillators (Figure 2b-e)^20,21^.

The absorption spectra of aSyn in the two salts were deconvoluted and the absorption coefficient of the solvation shell surrounding the protein was calculated for a range of different supposed shell sizes and compared to bulk absorption. In bulk, a solution containing CsI absorbs 2.66 times as much as one containing NaCl (flat black line, Figure 2f). When taking into account that the size of the solvation shell around the protein, which includes some ions, may depend on the salts, and especially the anions (calculated by AIMD in Supplementary Figure 5), the solvated protein in a solution containing CsI is predicted to absorb between 2.7 and 2.9 times as much as one in a solution containing NaCl in the largest mathematically possible solvation shell, i.e. in the case that solvation shells take up all available volume. It is found that the larger the assumed solvation shell, the larger its absorption coefficient, while still being lower than bulk absorption. The absorption in the solvation shell is thus directly influenced by the interaction of the protein and salts and cannot be explained by the different absorption of hydrated salt ions only. The absolute difference in absorption upon adding aSyn to a salt solution is lower in NaCl than in CsI, showing solute mobility in NaCl is lower than CsI.

We then investigated the absorption of solvation shell without the influence of the protein in the salt solutions in a second deconvolution step. As mentioned previously, the protein and shell absorb less THz radiation than the water-salt solution they replace (Figure 2b-e). To investigate the absorption of the solvation shell in the presence of the salts we first defined the parameters for salt-independent absorption, where the aSyn molar absorption coefficient (*ε*_P_) is not influenced by the hydration with different salts. As *ε*_P_ is not known, the salt-independent absorption coefficient of the solvation shell was calculated based upon protein absorption defined between the upper boundary of 25 cm^−1^M^−1^, which is comparable to the absorption of aSyn in solid CsI at room temperature, and the lower boundary value, 1 cm^−1^M^−1^, around the limit of detection. Between the two boundaries the ratio of the absorption of the solvent in the bulk and shell, independent of salt, is the same (Figure 2g).

We subsequently investigated the influence of the salt on the solvation shell, independent of the protein. The shell containing NaCl absorbs less than the bulk above *ε*_P_ = 5 cm^−1^M^−1^, whereas the shell containing CsI only absorbs less than the bulk above *ε*_P_ = 15 cm^−1^M^−1^ (Figure 2g). If *ε_p_* is higher than 15 cm^−1^M^−1^, both shells absorb less than the bulk, but the one containing NaCl even less so than the one including CsI. This shows that across a physiological range of protein absorption, ion and water mobility in the vicinity of aSyn are increased in CsI compared to NaCl. This is clearly observed at *ε*_P_ = 10 cm^−1^M^−1^ (shown by the circles in Figure 2g), an intermediate protein absorption coefficient, where the shell absorption in NaCl is reduced while increased in CsI. Our THz-TDS measurements have shown that adding the protein disturbs the interaction between water molecules and salt ions, and depends on the salt ion. This can result in an increase or decrease of mobility in the shell compared to the bulk.

### aSyn structure is similar when bound to Na^+^ and Cl^-^ as Cs^+^ and I^-^

We next investigated the possibility of structural differences of aSyn in the presence of NaCl and CsI to determine whether the difference in aggregation rate could instead be due to a direct interaction of NaCl or CsI with aSyn. Native nano-electrospray ionisation mass spectrometry (nano-ESI-MS) data show that aSyn binds a maximum of three Na^+^ and five Cs^+^ ions at a 1:50 ratio (20 μM aSyn: 1 mM salt) (Figure 3a) (data for 1:250 ratio is presented in the Supplementary Figure 7 and discussed in Supplementary Note 2). Binding of the counter anion I^-^ and Cl^-^ is not observed.

**Figure 3.**
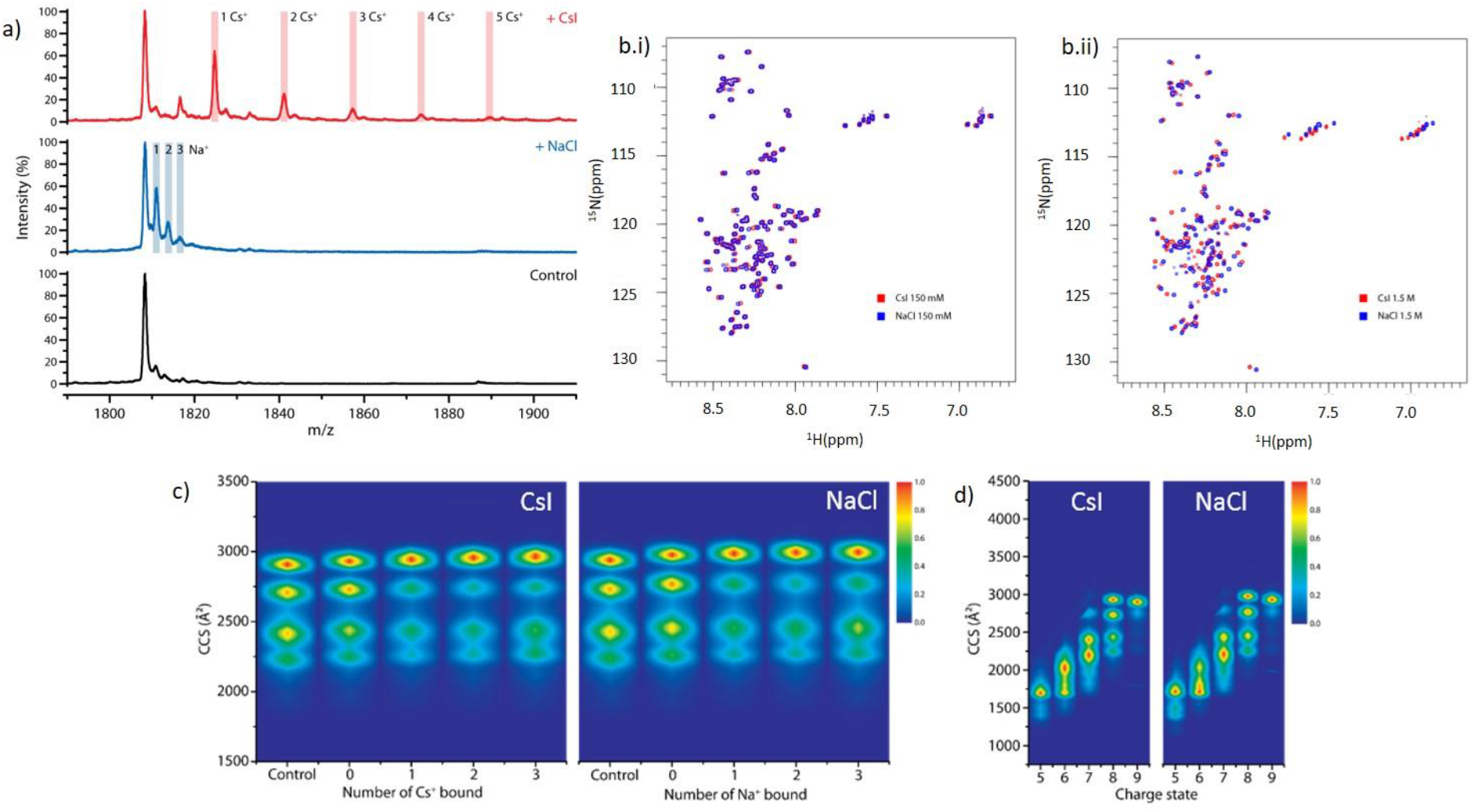
aSyn binds more Cs^+^ than Na^+^ which does not grossly affect aSyn conformation. The mass spectrum of (a) native aSyn (Control, black) is shown in the 8+ charge state region, and in the presence of a 1:50 ratio (20 μM aSyn: 1 mM salt) we observe aSyn bound to three Na^+^ (+NaCl, blue) and to five Cs^+^ (+CsI, red). (b) 2D ^1^H-^15^N HSQC peak spectrum of aSyn containing (b, i) 150 mM CsI (red) in 5% D_2_O, 95% H_2_O (vol/vol) was overlaid with aSyn containing 150 mM NaCl (blue) in 5% D_2_O, 95% H_2_O (vol/vol). (b, ii) aSyn with 1.5 M NaCl (red) (vol/vol) was overlaid with aSyn containing 1.5 M CsI (blue). Gross shift perturbations are only observed across the protein sequence under very high (1.5 M) salt concentrations. (c) Heat maps of the aSyn conformations detected for the 8+ charge state of aSyn without salt present and in the presence of 1 mM NaCl/CsI in a 1:50 protein:salt ratio, and displayed in the absence of ions (Control), in the presence of ions but not binding (0), and as a function of the number of cations bound (1, 2, 3). (d) Heat maps of the conformations of aSyn detected at different charge states (5+ to 9+) in the presence of 1 mM NaCl/CsI using a 1:50 protein:salt ratio.

^15^N-labelled aSyn was then measured by 2D ^15^N HSQC NMR spectroscopy to investigate structural changes in both 150 mM and 1.5 M CsI and NaCl solutions. The chemical shift spectrum of aSyn showed few chemical shift changes of residues in 150 mM salt solutions (Figure 3bi), however large chemical shift changes were observed across all regions of aSyn in 1.5 M salt solutions (Figure 3bii). Chemical shifts across all regions of aSyn suggest there are no specific binding regions for the ions, but that salt binding at very high, non-physiological concentrations may induce structural differences. However, the chemical shift changes observed by NMR display an ensemble measurement, likely representing the average of all states of aSyn in the salt solutions. As an IDP aSyn resides in many transient conformations, therefore we cannot clearly determine whether there are shifts in the distribution of aSyn conformations in the different salt solutions using this method.

We hence used nano-ESI-ion mobility-MS (nano-ESI-IM-MS) to investigate potential changes to the distribution of aSyn conformations when bound to the salt ions. In the ion mobility experiment, the amount of gas-phase collisions, and therefore the drift time, is directly related to the rotationally averaged extendedness of the protein ion^22^. Using the 8+ charge state of aSyn in the absence of salt, we identified four main co-existing conformations in the gas phase (Figure 3c), as previously reported^23^. The choice of charge state represented is discussed in Supplementary Note 3. The larger the collision cross sections (CCS), the more extended the protein structures. The binding of ions induced a shift favouring conformations with higher CCS values (Figure 3c). There were no further differences in the distribution of the conformational ensembles of aSyn in either salt with increasing numbers of ions bound or with increasing salt concentrations (Figure 3c, Supplementary Figure 8). Overall, the aSyn conformational space did not extend or compact drastically, as has been observed for the binding of some small molecule drugs to aSyn^23^, suggesting that the binding of these monovalent ions is non-specific, similar to what we observe by NMR and what we have observed with other monovalent ions^24^. At lower charge states, particularly 5+, 6+ and 7+, we observe a slight difference in the intensity distribution of aSyn conformations in CsI compared to NaCl containing solutions (Figure 3d). This suggests that there could be differences in the distribution of the structural ensemble of aSyn in the two salt solutions, however these charge states may represent conformations that are influenced by the gas phase. Both NMR and nano-ESI-IM-MS data suggest there are no gross differences in the conformation of aSyn in the presence of CsI or NaCl, but possibly the current resolution of these methods do not allow us to fully determine different structures in these ensemble solutions.

### *In vitro* aSyn is more mobile in CsI than in NaCl

We finally examined the mechanism of altered solvent mobility on the aggregation propensity of aSyn. MD simulations indicated that the altered water mobility in the solvation shell and the mobility of the aSyn_72-78_ peptide backbone were inextricably linked, and the mobility of the aSyn_72-78_ peptide chain in CsI solution was reduced compared to in NaCl. We therefore examined the rate of the conformational rearrangement of aSyn in the two salts using ^15^N HSQC NMR spectroscopy and THz-TDS. Both techniques showed that aSyn in NaCl a had reduced mobility compared to aSyn in CsI. The mobility of ^15^N-labelled aSyn was expressed through the signal intensity of the individual aSyn residues, the lower signal intensity of aSyn in the 1.5 M CsI solution is related to the protein being more mobile in CsI on a timescale that leads to a reduction in signal intensity compared to the NaCl solution (Figure 4a). At the lower salt concentration at 150 mM the difference in mobility was smaller, but still observed (Supplementary Figure 9). Furthermore, we observe that most of the protein sequence is influenced by the presence of NaCl and CsI as there are no specific binding sites or regions for the ions present, which may lead to more localised intensity changes, and region-specific peak shifts in the spectra for which none have been observed (Figure 4b). The N-terminal residues 1-20 were less influenced by the salt ions and were more similar in intensity.

**Figure 4.**
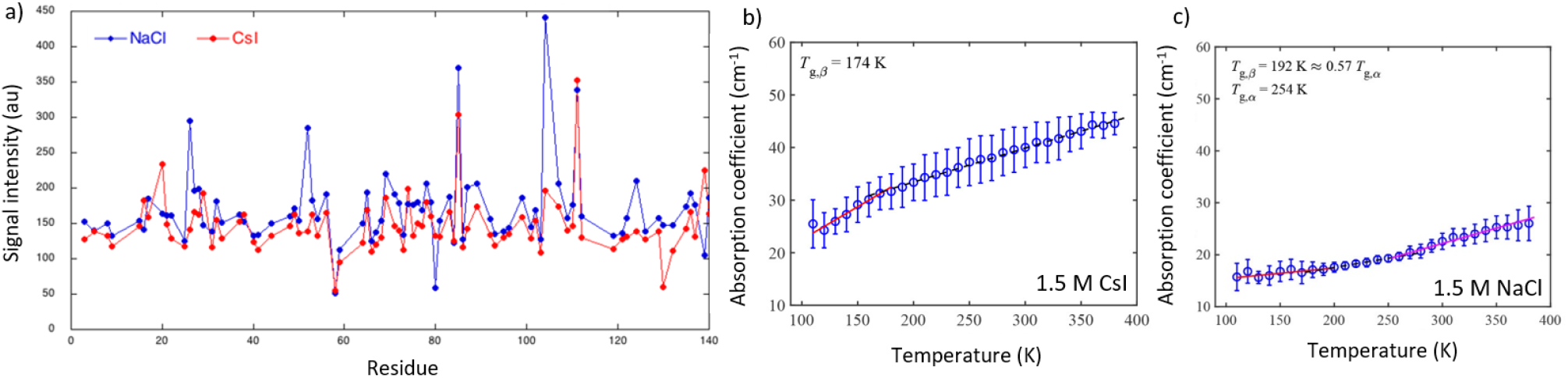
NMR and THz-TDS show that aSyn is more mobile in CsI compared to NaCl. (a) HSQC NMR spectroscopy was used to measure the intensity of 150 μM ^1^H and ^15^N-labelled aSyn in 95% H_2_O, 5% D_2_O (vol/vol) containing 1.5 M CsI (red) and NaCl (blue). The signal intensity for aSyn is displayed for each salt with 86% residue coverage. Each residue covered is represented by a dot. aSyn samples containing CsI had an overall lower intensity across most of the protein sequence. The mean terahertz absorption coefficient as a function of temperature at 1 THz is shown for (b) aSyn and 1.5 M CsI and (c) aSyn and 1.5 M NaCl. Lines indicate the different linear fits of the respective regions. Error bars represent the standard deviation of 3 measurements.

Temperature ramping with THz-TDS showed that a solid state sample of aSyn and CsI had a secondary glass transition temperature at *T*_g,*β*_ = 174 K while for aSyn and NaCl *T*_g,*β*_ = 192 K. The glass transition at *T*_g,*β*_ is associated with the onset of local mobility of the sample^25^ and aSyn samples containing CsI become mobile at a lower temperature than aSyn samples containing NaCl (Figure 4b, Table 4). Furthermore, the gradient of the slope at temperatures below *T*_g,*β*_, which represents a measure of inherent molecular mobility, is steeper for aSyn and CsI (*m* = 0.12 cm^−1^ K^−1^) than aSyn and NaCl (*m* = 0.02 cm^−1^ K^−1^) indicating that aSyn with CsI is more mobile than for aSyn with NaCl. The aSyn and NaCl sample displays a second transition temperature, *T*_g,*α*_ = 254 K, which refers to large scale mobility attributed to cooperative motions of the sample (Figure 4c, Table 4). No distinct *T*_g,*α*_ was observed for the aSyn sample containing CsI indicating that there is sufficiently high mobility already present at lower temperatures, hence the cooperative motions gradually emerge at temperatures above *T*_g,*β*_ instead of being associated with a defined transition point. The THz-TDS data are in agreement with our NMR data which show that aSyn samples containing CsI are more mobile and able to reconfigure than aSyn samples containing NaCl.

**Table 4.** Gradient, *m*, of the linear fit (*y* = *mx+c*) and respective glass transition temperatures determined using THz-TDS

| Sample | Region 1<br>($\text{cm}^{-1} \text{K}^{-1}$) | Region 2<br>($\text{cm}^{-1} \text{K}^{-1}$) | Region 3<br>($\text{cm}^{-1} \text{K}^{-1}$) | $T_{g,\beta}$<br>(K) | $T_{g,\alpha}$<br>(K) |
| --- | --- | --- | --- | --- | --- |
| aSyn + CsI | $0.12 \pm 0.01$ | $0.067 \pm 0.001$ | - | 174 | - |
| aSyn + NaCl | $0.02 \pm 0.01$ | $0.035 \pm 0.001$ | $0.059 \pm 0.01$ | 192 | 254 |

## Discussion

The influence of ions on the mobility of water has been well studied, yet the effect of water mobility on the propensity of proteins to misfold is still not elucidated and in particular, not in connection with IDPs and amyloid fibril formation. Here, we show that ions can influence the mobility of bulk water, water in the solvation shell and protein mobility, and that the dynamics of the aqueous phase governs aggregation rates. The trend for increased aSyn aggregation rate in NaCl compared to CsI is observed in both H_2_O and in D_2_O, yet the aggregation rates of aSyn were faster in D_2_O. This suggests the direct effect on aggregation comes from the solvent and that the ions can influence the solvent. The presence of deuterium bonds, which are stronger and shorter than hydrogen bonds^26,27^, may increase aggregation propensities of proteins^28–32^. We directly observe that the presence of CsI leads to increased water mobility, both in bulk and in the protein solvation shell, in comparison to NaCl. An increase in absorption as measured with THz-TDS directly relates to an increased change in dipole moment and therefore ion and protein mobility which are inextricably linked to the mobility of surrounding water molecules.

Although direct ion binding has been proposed to influence aSyn aggregation rates, the ion binding strength does not correlate with aggregation rates observed^33^, suggesting that the Hofmeister series may not be the only explanation for why these ions either decrease or increase aSyn aggregation kinetics. Furthermore, we can exclude the Debye-Hückel effect as both NaCl and CsI are monovalent; if such a charge screening effect was dominant, a similar effect on the aggregation kinetics of aSyn should have been observed. Structural alterations to the dynamic ensemble of aSyn conformations by NaCl and CsI, which may favour aggregation prone conformations, cannot be ruled out. Although we observed no gross differences in the structures of aSyn by NMR and MS in the presence of NaCl and CsI, these techniques may not be sensitive enough on the timescale needed to identify differences in transient dynamic interactions within the monomer structures in solution. Yet, these dynamic interactions govern whether a protein remains monomeric or misfolds into conformations that can aggregate. The surrounding solvent dictates the time scale for forming and maintaining these conformations.

IDPs rely on their ability to be highly dynamic and flexible to probe different conformational space allowing maintenance of their solubility and function. When the reconfiguration rates of the protein backbone are retarded this can lead to aggregation^17,34–38^. For protein association and aggregation to occur, the proteins must firstly be in an aggregation prone conformation, and secondly must be stable for long enough for interactions to occur. Our data support a mechanism whereby the pathway to oligomerisation and aggregation is determined by the intramolecular diffusion rate of the protein, which we show is determined by the mobility of, and intermolecular interactions with, the surrounding water, which in turn is modulated by of ions present (Figure 5). These data may have important implications for aSyn localised within certain environments either inside or outside of a cell, where ion concentrations can differ greatly^39,40^. The presence of ions during the formation of the yeast prion protein oligomers, but not during elongation, can influence fibril polymorphism and is directly linked to pathology^41,42^. Therefore, interesting questions arise regarding cell specific or age dependent accumulation of certain ions or metabolites in the intracellular aqueous environment that could alter water mobility and influence aSyn strain polymorphism and disease outcome.

**Figure 5.**
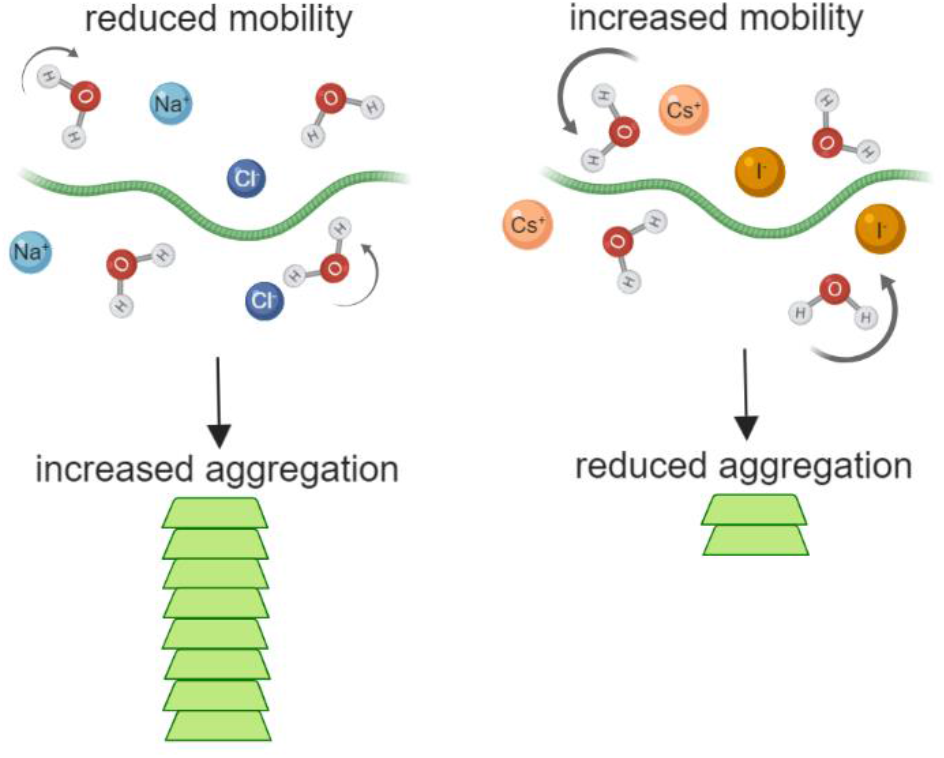
Aggregation kinetics of aSyn are dependent on water mobility which is strongly affected by the presence of ions. Presence of Na^+^ (light blue) and Cl^-^ (dark blue) lead to reduced mobility of water (H_2_O) and aSyn monomers (green protein), allowing the formation of stable intermolecular bonds between two adjacent monomers which can lead to aggregation into aSyn amyloid fibrils (stacked green protein). Presence of Cs^+^ (light orange) and I^-^ (dark orange) lead to increased mobility of water and aSyn monomer, which decreases the likelihood of two monomers being stable enough to permit intermolecular interactions and thus results in reduced aggregation.

## Methods and Materials

### Purification of aSyn

Human wild-type (WT) alpha-synuclein was expressed using plasmid pT7-7. The plasmid was heat shocked into *Escherichia coli* One Shot^®^ BL21 STAR™ (DE3) (Invitrogen, Thermo Fisher Scientific, Cheshire, UK) and purified as previously described^43^. Recombinant aSyn was purified using ion exchange chromatography (IEX) in buffer A (10 mM Tris, 1 mM EDTA pH 8) against a linear gradient of buffer B (10 mM Tris, 1 mM EDTA, 0.5 M NaCl pH 8) on a HiPrep Q FF 16/10 anion exchange column (GE Healthcare, Uppsala, Sweden). aSyn was then dialysed into buffer C (1 M (NH_4_)_2_SO_4_, 50 mM Bis-Tris pH 7) and further purified on a HiPrep Phenyl FF 16/10 (High Sub) hydrophobic interaction chromatography (HIC) column (GE Healthcare) and eluted against buffer D (50 mM Bis-Tris pH 7). Purification was performed on an ÄKTA Pure (GE Healthcare). aSyn was extensively dialysed against H_2_O and used immediately for experiments or dialysed against 20 mM Tris pH 7.2 and concentrated using 10 k MWCO amicon centrifugal filtration devices (Merck KGaA, Darmstadt, Germany) and stored at −80 °C until use. aSyn in Tris was buffer exchanged into freshly prepared NaCl and CsI solutions (pH 5.4) before experiments using PD10 dialysis columns (GE Healthcare). Protein concentration was determined from the absorbance measurement at 280 nm on a Nanovue spectrometer using the extinction coefficient for aSyn of 5960 M^−1^cm^−1^.

Protein purity was analysed using analytical reversed phase chromatography (aRP). Each purification batch was analysed using a Discovery BIO Wide Pore C18 column, 15cm × 4.6mm, 5μm, column with a guard cartridge (Supelco by Sigma-Aldrich, St. Louis, MO, USA) with a gradient of 95 % to 5 % H_2_O + 0.1% trifluroacetic acid (TFA) or acetic acid and acetonitrile + 0.1% TFA or acetic acid at a flow-rate of 1 mL/min. The elution profile was monitored by UV absorption at 220 nm and 280 nm on an Agilent 1260 Infinity HPLC system (Agilent Technologies LDA, Santa Clara, USA) equipped with an autosampler and a diode-array detector. Protein purity fell between 88 - 95 % dependent on batch (Supplementary Figure 10).

### Thioflavin-T based assays

aSyn samples were buffer exchanged into 150 mM NaCl, 1.5M NaCl, 150 mM CsI or 1.5 M CsI in H_2_O or D_2_O prior to performing experiments using PD10 desalting columns (GE Healthcare). 10 μM freshly made ThT (abcam, Cambridge, UK) was added to 50 μL of 50 μM aSyn. All samples were loaded onto nonbinding, clear bottom, 365-well black plates (Greiner Bio-One GmbH, Kremsmünster, Austria). The plates were sealed with a SILVERseal aluminium microplate sealer (Grenier Bio-One GmbH). Fluorescence measurements were taken using a FLUOstar Omega plate reader (BMG LABTECH GmbH, Ortenberg, Germany). The plates were incubated at 37 °C with orbital shaking at 300 rpm for five minutes before each read every hour. Excitation was set at 440 nm with 20 flashes and the ThT fluorescence intensity was measured at 480 nm emission with a 1300 gain setting. ThT assays were repeated at least three times using at least three wells for each condition. Data were normalised to the well with the maximum fluorescence intensity for each plate and the average was calculated for all experiments. Data are displayed with the rolling average from three experiments calculated using the program R (https://www.r-project.org/) with a 0.5 span. A linear trend line fitted along the exponential phase region of the ThT fluorescence curve was used to calculate the lag time (t_lag_) using equation 1.

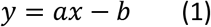

### Determination of remaining monomer concentration of aSyn after ThT-based assays using analytical size exclusion chromatography

To calculate the remaining aSyn monomer concentration in each well after ThT-based assays size exclusion chromatography performed on a high-pressure liquid chromatography (SEC) system was used. The contents of each well after the ThT-based assay were centrifuged at 21k × g for 20 minutes and the supernatant from each well was added to individual aliquots in the autosampler of the Agilent 1260 Infinity HPLC system (Agilent Technologies). 35 μL of each sample was injected onto an Advance Bio SEC column, 7.8 × 300 mm 130Å (Agilent Technologies) in 20 mM Tris pH 7.2 at 1 mL/min flow-rate. The elution profile was monitored by UV absorption at 220 and 280 nm. A calibration curve of known concentrations of aSyn was used to calculate the remaining monomer concentration of aSyn in each well. Six wells from two experiments were analysed for remaining monomer concentrations, the average value of each measurement is presented including the standard error of the mean (SEM).

### Atomic Force Microscopy

The contents of wells from the ThT-based assays were centrifuged for 20 minutes at 21 k × g. 40 μL of supernatant was removed to leave 10 μL and remaining fibrils. The fibrils were resuspended and incubated on a freshly cleaved mica surface for 20 min. The mica was washed three times in 18.2 Ω dH_2_O to remove lose protein. Images were acquired in dH_2_O using tapping mode on a BioScope Resolve (Bruker GmbH, Karlsruhe, Germany) using ‘ScanAsyst-Fluid+’ probes. 256 lines were acquired at a scan rate of 0.966 Hz per image with a field of view of 2-5 μm and for at least six fields of view. Images were adjusted for contrast and exported from NanoScope Analysis 8.2 software (Bruker).

### Small Angle Neutron Scattering

Small-angle neutron scattering (SANS) measurements were performed on the SANS2D instrument at the ISIS Neutron AND Muon Source (STFC Rutherford Appleton Laboratory, Didcot, UK). 6.25 mg of freeze-dried aSyn was dissolved in a salt solution containing 1.5 M NaCl or CsI in Milli-Q Ω18.2 water or pure deuterated water to give a final concentration of 434 μM. The protein solution was left stirring in a cooling cabinet for 0.5 h in H_2_O salt solutions, whereas for D_2_O salt solutions it was left for 1.5 h to allow for sufficient hydrogen/deuterium exchange. Samples were then loaded into quartz circular cells of 1 mm (H_2_O samples) and 2 mm (D_2_O samples) pathlength and measurements were made at room temperature. The protein solutions were also measured 15 to 19 hours after preparation.

An incident beam of 12 mm diameter, a wavelength range of 1.75–16.5 Å, and a setup of L_1_ = 4 m; L_2_ = 4 m, was used resulting in an effective range of wave vector in equation 2,

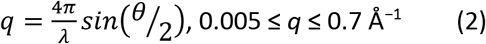

where *λ* = the neutron wavelength and *θ* = the scattering angle. Raw scattering data are corrected for sample transmission, detector efficiency and solvent background scattering (as described in detail in^44^) using the Mantid Software, and then converted to absolute scattering cross section (*I*(*q*) / cm^−1^) using the scattering from a standard sample (comprising a solid blend of hydrogenous and perdeuterated polystyrene) in accordance with established procedures^45^.

Modelling of the data was performed in SASView (http://www.sasview.org), using the Guinier–Porod model^46^. The scattering intensity, *I*(*q*), is derived from independent contributions of the Guinier form in equation 3,

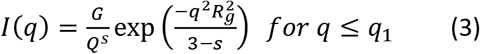

and the Porod form, in equation 4

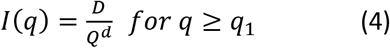

where *q* is the the scattering variable, *R_g_* is the the radius of gyration, *d* is the the Porod exponent, and *G* and *D* are the Guinier and Porod scale factors respectively. A dimensionality parameter (3 – *s*) is included in the Guinier form factor to help define non-spherical objects where *s* = 0 represents spheres or globules, *s* = 1 represents cylinders or rods and *s* = 2 represents lamellae or platelets.

### Ab Initio Molecular Dynamics (AIMD)

The CP2k software package was used for all AIMD simulations, which incorporated three-dimensional periodic boundary conditions^47,48^. The simulations made use of the Perdew-Burke-Ernzerhof (PBE) density functional^49^ coupled with the dispersion correction of Grimme^50,51^. The electronic wave functions were represented using the double-zeta DZVP basis set^47^. Simulations were performed within the canonical ensemble (NVT), with the temperature maintained at 300 K using a Nose-Hoover chain thermostat^52–54^. The crystal structure of seven amino acids, TGVGAGA residues 72-78 of aSyn, was taken from the protein database for simulations^55^. The initial model was generated by fully solvating the aSyn peptide with water molecules explicitly in a 125 nm^3^ simulation box. Ions were introduced to a concentration of 1.5 M to match the experiment. The simulations were equilibrated for 5 ps prior to performing the production MD runs over a 45 ps trajectory (50 ps total), with a time step of 1.0 fs.

### Terahertz spectroscopy in liquid

aSyn was dialysed extensively against H_2_O to remove salts after purification. The samples were snap frozen in liquid nitrogen and lyophilised using a LyoQuest 85 freeze-dryer (Telstar, Spain). The aSyn samples were resuspended at a concentration of 691.56 μM (10 mg/mL). 10 mM Tris pH 7.2 was added to the samples to aid reconstitution. Samples were reconstituted in the salts and sonicated for 10 s on and 10 s off for three times before THz measurements. The liquid was injected into a liquid cell with a path length of 100 μm. Reference measurements of buffer were performed using the same liquid cell. THz-TDS spectra were acquired using a commercial TeraPulse 4000 instrument with a spectral range of 0.2-2.7 THz (TeraView, Cambridge, UK). The temperature was kept constant at 294 K. The absorption coefficient of the liquid samples was calculated in the same way as that of solid samples. Measurements were repeated at least 5 times. Buffer and salt solutions were measured in 0.25 M increments for NaCl concentrations of 0.5 to 4 M, and for CsI concentrations of 0.25 to 2.5. aSyn in NaCl was measured at 2 M, as aSyn did not reconstitute successfully in concentrations below 2 M, and aSyn in CsI was measured at 1.25, 1.5, and 2 M. All spectra were subsequently divided by the salt concentration to obtain the molar absorption coefficient *ε*. *ε* was then fitted over frequencies with a linear function for samples containing NaCl. In samples containing CsI a spectral feature appeared at 0.7 THz and *ε* was therefore fitted with the sum of a power law and a Lorentzian to incorporate the spectral features. [REFs]

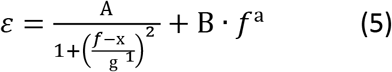

Where A is the peak intensity, g the half width at half maximum, x1 the centre frequency of the peak, f the frequency, and a and B are power law parameters. An offset was not observed in any measurement for which reason no absolute term is present.

The THz-TDS spectra of liquid aSyn in the two salts were deconvoluted to investigate the effects of the salt ions on the solvation shell. The solvation shell size of the single ions was based on the results obtained from the AIMD simulations. Because the absorption coefficients of the salts without protein present are known, absorption coefficients of the protein and its solvation shell can be calculated for different estimates of solvation shell sizes.

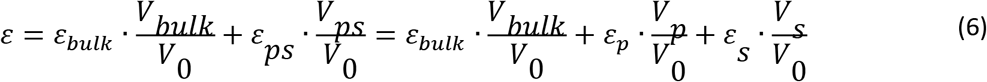

### Purification of aSyn for NMR analysis

*E. coli* were grown in isotope-enriched M9 minimal medium containing ^15^N ammonium chloride similar our previous protocol^56^. To isolate expressed aSyn the cell pellets were resuspended in lysis buffer (10mM Tris-HCl pH 8, 1mM EDTA and EDTA-free complete protease inhibitor cocktail tablets (Roche, Basel, Switzerland), 0.2 mM phenylmethylsulfonyl fluoride (PMSF) and Pepstatin A) and lysed by sonication. The cell lysate was centrifuged at 22k × g for 30 min to remove cell debris and the supernatant was then heated for 20 min at 90 °C to precipitate the heat-sensitive proteins and subsequently centrifuged at 22k × g. Streptomycin sulfate 10mg/ml was added to the supernatant to precipitate DNA. The mixture was stirred for 15 min followed by centrifugation at 22k × g, then repeated. Ammonium sulfate 360 mg/ml was added to the supernatant precipitate the protein aSyn. The solution was stirred for 30 min and centrifuged again at 22k × g. The resulting pellet was resuspended in 25mM Tris-HCl, pH 7.7 and dialyzed overnight. The protein was purified by IEX on a HiPrep Q FF anion exchange column (GE Healthcare) and then further purified by SEC on a HiLoad 16/60 Superdex 75 prep grade column (GE Healthcare). All the fractions containing the monomeric protein were pooled together and concentrated using amicon 10 k MWCO centrifugal filtration devices (Merck). Protein purity was determined by aRP to be 88.6% (Supplementary Figure 8). aSyn was buffer exchanged into 5% D_2_O and 95% H_2_O using PD10 desalting columns (GE Healthcare). CsI and NaCl were added to a final concentration of 150 mM and 1.5 M just before performing the experiments.

### Nuclear Magnetic Resonance Spectroscopy

NMR spectra were recorded as 2D ^15^N HSQC at 298 K on a Bruker AV800 spectrometer (800 MHz ^1^H) equipped with a 5 mm TXI HCN/z cryoprobe. Increase in salt concentration from 150 mM to 1.5 M resulted in an intrinsic signal intensity loss of 2.25x for NaCl and CsI, due to increased lossiness of the sample. Accordingly, spectra recorded at 1.5 M salt were multiplied by a factor of 2.25x prior to intensity analysis. Experiments were recorded with 2 scans per free induction decay with 150 and 1024 complex pairs in ^15^N and ^1^H, respectively.

### Native nano-electrospray ionization mass spectrometry (nano-ESI-MS) and ion mobility (IM)

aSyn was buffer exchanged into 20 mM ammonium acetate (Sigma Aldrich, St. Louis, MO, USA) pH 7 using PD 10 columns (GE Healthcare) and diluted to a final concentration of 20 μM. NaCl (Acros Organics, New Jersey, USA) or CsI (Sigma Aldrich, St. Louis, MO, USA) were dissolved in water and added to the sample with a final concentration between 80 μM and 5 mM, which corresponds to a 1:4, 1:50 and 1:250 aSyn:Na/Cs ratio. The samples were incubated for ten minutes at room temperature before analysis. A Synapt G2 HDMS (Waters, Manchester, UK) was used to perform the nano-ESI (ion mobility-) mass spectrometry (nano-ESI-IM-MS) measurements. The results were analysed using Masslynx version 4.1 (Waters, Manchester, UK). Infusion of the samples into the mass spectrometer was performed using home-made gold-coated borosilicate capillaries. The main instrumental settings were: capillary voltage 1.5-1.8 kV; sampling cone 25 V; extraction cone 1 V; trap CE 4 V; transfer CE 0 V; trap bias 40 V. Gas pressures used throughout the instrument were: source 2.75 mbar; trap cell 2.3 × 10^−2^ mbar; IM cell 3.0 mbar; transfer cell 2.5 × 10^−2^ mbar. In mass spectra of the aSyn + CsI sample there are low intensity Na^+^ adducts remaining bound in spite of buffer exchange.

### Terahertz time domain spectroscopy (THz-TDS) of solid samples

aSyn (2.2 mg/ml) was buffer exchanged into 1.5 M NaCl or 1.5 M CsI in H_2_O using PD10 desalting columns. The samples were snap frozen in liquid nitrogen and lyophilised using a LyoQuest 85 freeze-dryer (Telstar, Spain). The samples were prepared into pellets between 300-600 μm in thickness as outlined previously^57^. This sample was sandwiched between the two z-cut quartz windows and sealed in the sample holder. The THz-TDS spectra were acquired using a commercial TeraPulse 4000 instrument across a spectral range of 0.2-2.7 THz (TeraView, Cambridge, UK). The experiments were conducted over a range of temperatures (100-390 K) using a continuous flow cryostat with liquid nitrogen as the cryogen (Janis ST-100, Wilmington MA, USA) as outlined previously^57^. In order to calculate the absorption coefficient and the refractive index of the sample a modified method for extracting the optical constants from terahertz measurements based on the concept introduced by Duvillaret et al. was used^58,59^. The changes in sample dynamics were analysed by investigating the change in the absorption coefficient at a frequency of 1 THz and as a function of temperature. We have previously demonstrated that discontinuities in the temperature dependent absorption data in disordered materials can be used to highlight changes in the molecular dynamics and implemented a rigorous fitting routine based on statistical analysis to analyse the data outlined.

## Supporting Information

Data for purity of aSyn samples, analytical SEC to determine remaining monomer concentration after ThT-based kinetic assays, AFM data for the morphology of aSyn fibrils, additional SANS data analysis, additional nano-ESI-MS data, additional nano-ESI-IM-MS data, additional NMR data. Raw data is available at the University of Cambridge Repository. The SANS experiment at the ISIS Neutron and Muon Source was allocated under the beamtime XB1890203 (DOI: http://doi.org/10.5286/ISIS.E.RB1890203-1).

## Author Information

### Corresponding Author

Gabriele S. Kaminski Schierle

### Author Contributions

A.D.S. and G.S.K.S. conceived the project and designed experiments. A.D.S purified protein for all experiments, performed kinetic aggregation assays and AFM. N.M. performed SANS experiments, N.M and T.M analysed SANS data. MD simulations were performed by M.T.R. J.K. performed solution THz-TDS experiments. R.M. performed nano-ESI-MS and nano-ESI-IM-MS experiments. D.N. performed NMR experiments. T.S. performed solid THz-TDS experiments. All authors contributed to editing the manuscript and have given approval to the final version of the manuscript.

### Notes

The authors declare no competing financial interest.

## Acknowledgments

G.S.K.S. acknowledges funding from the Wellcome Trust (065807/Z/01/Z) (203249/Z/16/Z), the UK Medical Research Council (MRC) (MR/K02292X/1), Alzheimer Research UK (ARUK) (ARUK-PG013-14), Michael J Fox Foundation (16238) and Infinitus China Ltd. A.D.S. acknowledges Alzheimer Research UK for travel grants. M.T.R. and J.A.Z. acknowledge funding from EPSRC UK (EP/N022769/1). J.K. and J.A.Z. acknowledge funding from EPSRC (EP/S023046/1) and AstraZeneca. This work benefitted from SasView software, originally developed by the DANSE project under NSF award DMR-0520547.

## Supplementary Information

**Supplementary Figure 1.**
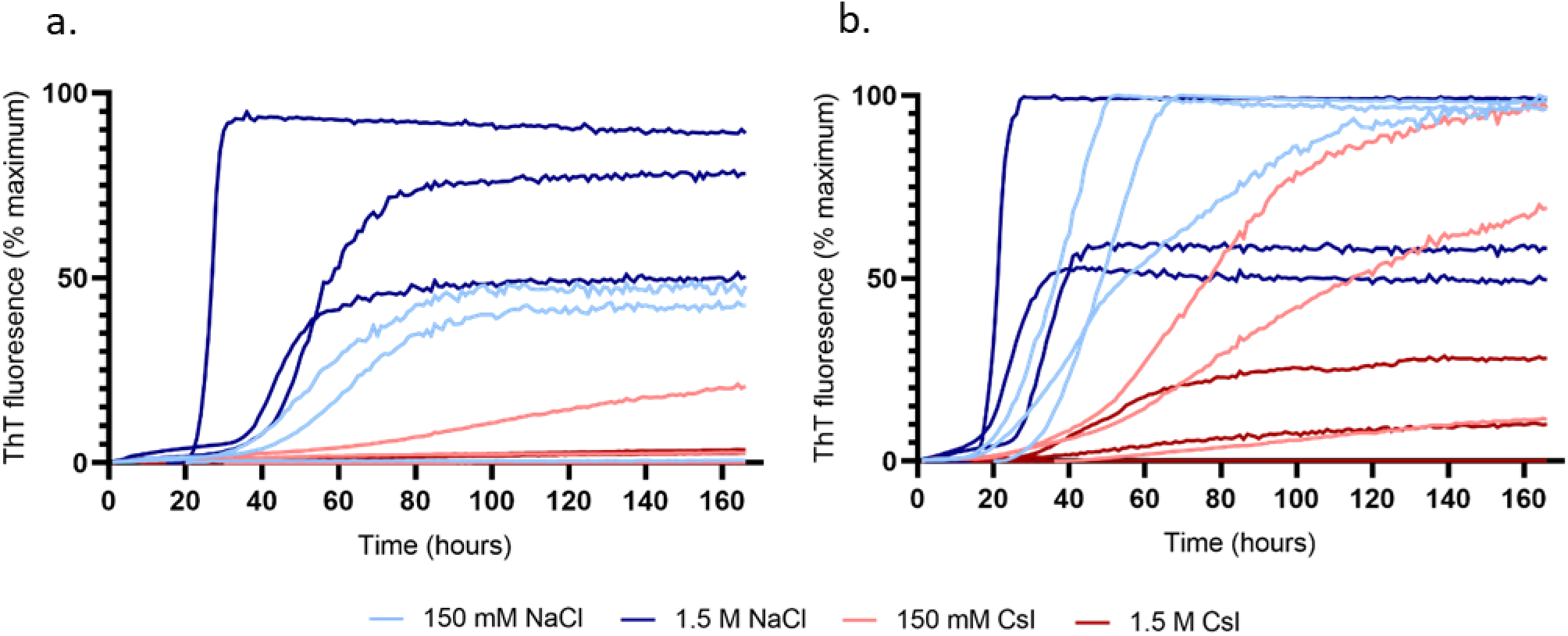
aSyn aggregation kinetics are enhanced in the presence of D_2_O and increasing concentrations of NaCl. aSyn aggregation kinetics were measured by ThT fluorescence; 50 μM aSyn was incubated with 10 μM ThT in a 384 well plate with continuous orbital shaking for 160 hours in the presence of (a) H_2_O and (b) D_2_O with 150 mM NaCl (red), 1.5 M NaCl (brown), 150 mM CsI (blue), 1.5 M CsI (navy) and plotted as % maximum ThT fluorescence over time. Increased NaCl concentrations accelerated aSyn aggregation, while increased CsI concentrations decelerated aSyn aggregation. The aggregation rate in D_2_O was enhanced compared to H_2_O. Three plate repeats are shown for each condition, the data presented are the average of four wells per condition.

**Supplementary Figure 2.**
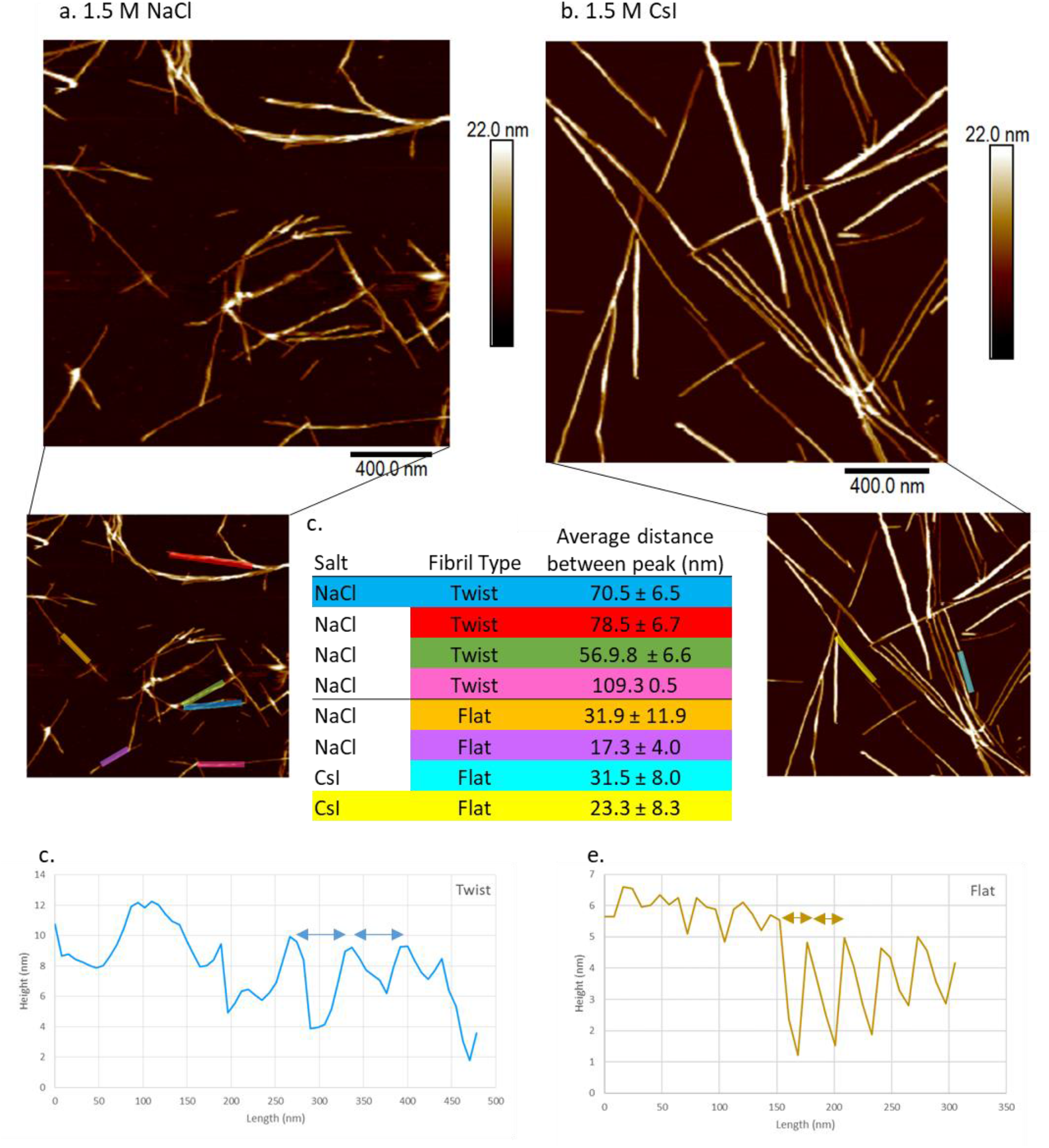
Identification of flat and twisted aSyn fibril polymorphs by AFM. aSyn fibrils formed during ThT-based assays were imaged on freshly cleaved mica and representative images are shown for different fibril polymorphs, ‘twisted’ containing a helical pitch and ‘flat’ with no visible helical pitch in (A.) 1.5 M NaCl in D_2_O and (B.) flat only 1.5 M CsI in D_2_O. Insets with coloured lines correspond to the colours in the table and show regions where fibril height was analysed. (C.) Table listing salt conditions, fibril type and distance measured between fibril peaks. Distances between peaks were calculated based on height profiles determined in the Nanoscope analysis software and are represented in (D.) for twisted fibrils with peak distances between 57-109 nm and (E.) flat fibrils with peak distances between 17 – 32 nm. The colours of the graph of peak heights in (D. + E.) correspond to the top blue twisted fibril and the bottom yellow flat fibril highlighted in table C and in the inserts of A and B.

**Supplementary Figure 3.**
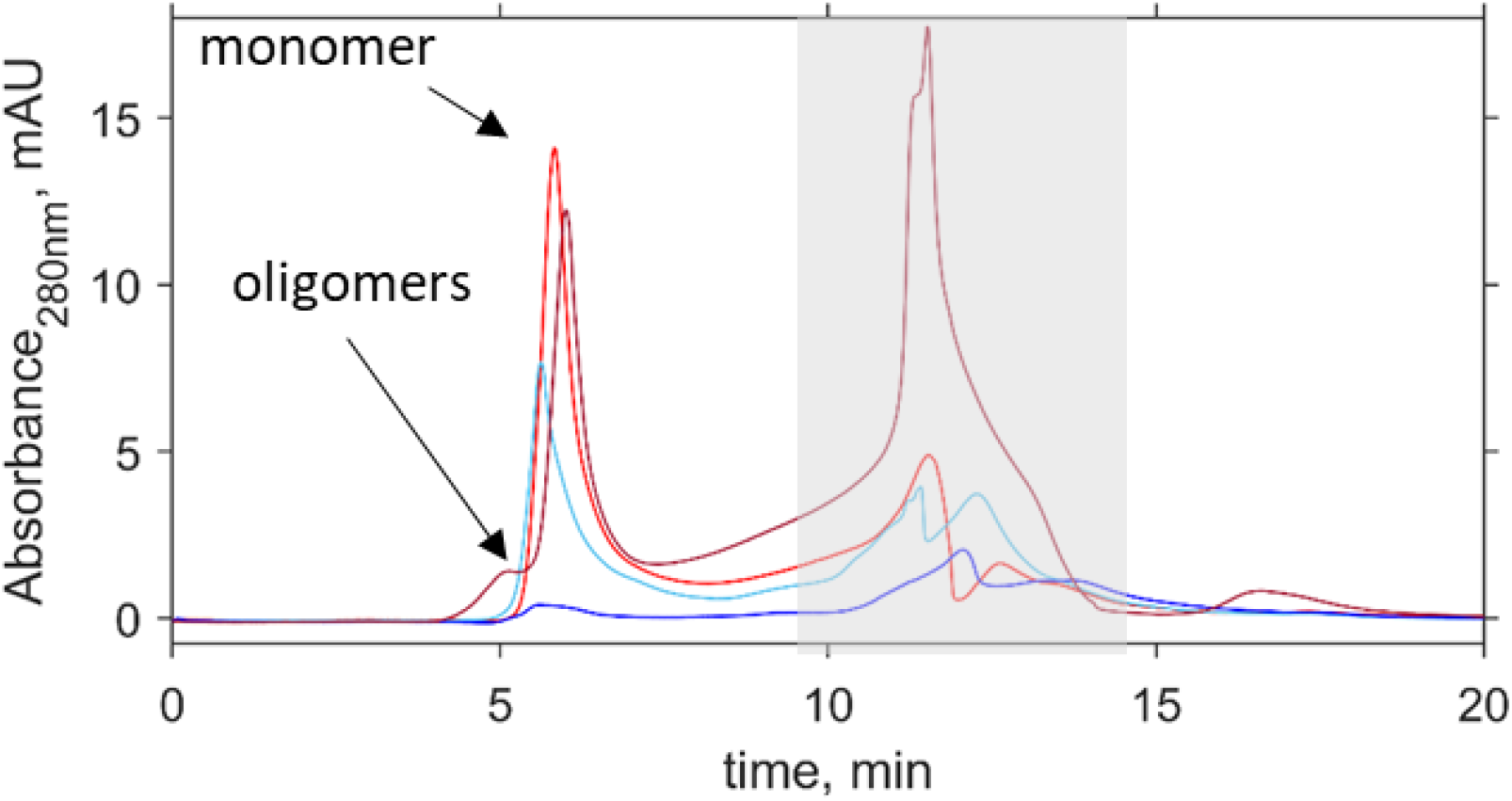
Representative analytical size exclusion chromatograph of remaining aSyn monomer after ThT-based assays. The content of each well after ThT-based assays was centrifuged and the remaining monomer analysed by analytical SEC-HPLC on an AdvanceBio 130Å column at a flow rate of 1 mL/min in 20 mM Tris pH 7.2 and monitored by absorbance at 280 nm. The area under the curve of the monomeric aSyn, which eluted ^~^ 5.2 minutes, was used to calculate the remaining monomer from known concentrations of aSyn. Representative chromatographs for aSyn in 150 mM NaCl (light blue, 1.5 M NaCl (dark blue), 150 mM CsI (red), 1.5 M CsI (dark red) are shown. Oligomeric species eluted before the monomer and can be detected in the 1.5 M CsI trace (dark red). Elution time for aSyn shifts slightly dependent on the salt and concentration present. The presence of salt is also detected in the grey region.

**Supplementary Figure 4.**
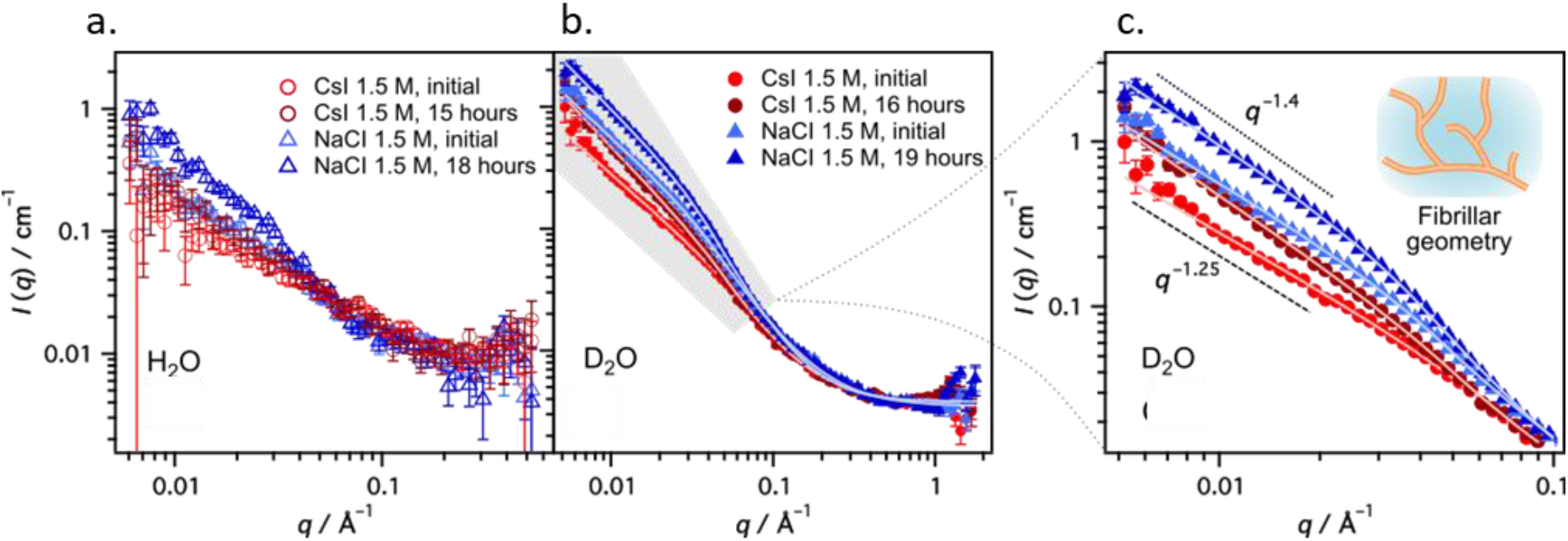
Model fits to the SANS data using the Guinier-Porod model. SANS data for solutions of 434 μM aSyn in either 1.5 M NaCl or CsI in H_2_O (a) and D_2_O (b). The aSyn in H_2_O samples had high scattering, therefore the model was fit to D_2_O data only. In c, solid symbols indicate experimental scattering data and solid lines represent the model fits. (g) Zoomed-in view of the low *q* region in b (shaded area) highlighting the differences in scattering intensity for each sample. Characteristic slopes are indicated with the data (q^−1.25^ and q^−1.4^), and the inset has been added to signify the fibrillar-type structure as determined by the modelling. Fitting values are shown in Supplementary Table 1.

**Supplementary Table 1.** Parameters of fitting SANS data presented in Figure 2 e,f. These results were obtained using a flexible cylinder model. The cylinder length fitted is not included as it is not detectable with the experimental q-range

| Solvent | Salt | Time | R (Å) | b (Å) | PDI of b | $\chi^2$ |
| --- | --- | --- | --- | --- | --- | --- |
| H<sub>2</sub>O | NaCl | Initial | 21 ± 3 | 199 ± 116 | 0.95 ± 0.36 | 0.63 |
|  | NaCl | 18 hours | 27 ± 4 | 116 ± 61 | 0.48 ± 0.18 | 0.8 |
|  | CsI | Initial | 20 ± 0.2 | 480 ± 42 | 0.11 ± 0.14 | 0.78 |
|  | CsI | 15 hours | 20 ± 0.9 | 468 ± 72 | 0.38 ± 0.17 | 0.8 |
| D<sub>2</sub>O | NaCl | Initial | 20 ± 0.1 | 103 ± 14 | 0.65 ± 0.26 | 3.2 |
|  | NaCl | 19 hours | 22 ± 1.1 | 100 ± 5.7 | 0.99 ± 0.11 | 2.93 |
|  | CsI | Initial | 20 ± 3.9 | 467 ± 111 | 0.91 ± 0.33 | 1.94 |
|  | CsI | 16 hours | 23 ± 4.2 | 409 ± 36 | 0.98 ± 0.38 | 2.65 |

**Supplementary Table 2.**
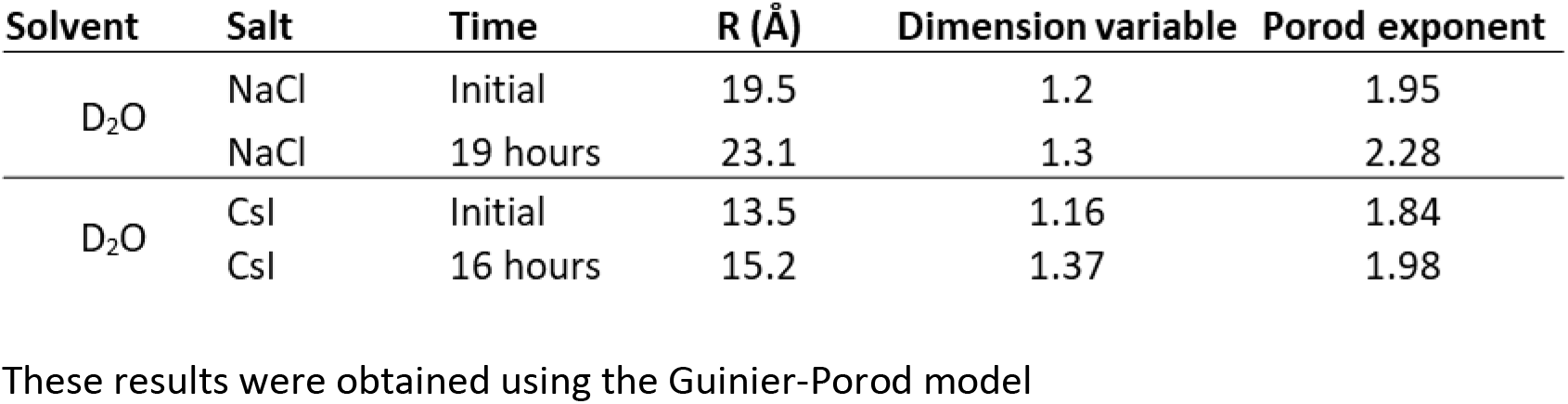
Parameters of fitting SANS data presented in Supplementary Figure 4

### Supplementary Note 1

#### Discussion of SANS data fitted with different models

The flexible cylinder model (Figure 1e,f, Table S1) gives a radius, R and, Kuhn length, b, that describes the stiffness of the fibril which can be related to shortness of fibril^60^. The fitting using the cylinder models to the scattering from aSyn in D_2_O show some discrepancy especially at high *q*-range, indicating that other populations (monomers or/and oligomers) might influence the scattering curves or that fibrils in D_2_O are not perfectly cylindrical. Instead we used a flexible cylinder and a sphere fitting (Figure 2g, Table 2 main text), to account for the contribution from the fibrils and monomers, respectively.

The Guinier-Porod model (Figure S4, Table S1) gives the radius, R, and a dimensionality parameter (3 – *s*) to help define non-spherical objects, where *s* = 0 represents spheres or globules, *s* = 1 represents cylinders or rods and *s* = 2 represents lamellae or platelets ^41^. Both fits give an average radius of ^~^20 Å, but the radius increases over time for all aSyn samples apart from aSyn in H_2_O with CsI. The Guinier-Porod model shows evidence of monomers in the high *q* region and the presence of flexible rod-like structures where s = 1.2-1.4 (Table S1). The higher slope of the aSyn in NaCl samples in D_2_O at low *q* indicates less rigid, i.e. longer aggregates, this is also observed in the cylinder model, where aSyn structures in NaCl are less rigid (Table 2).

**Supplementary Figure 5.**
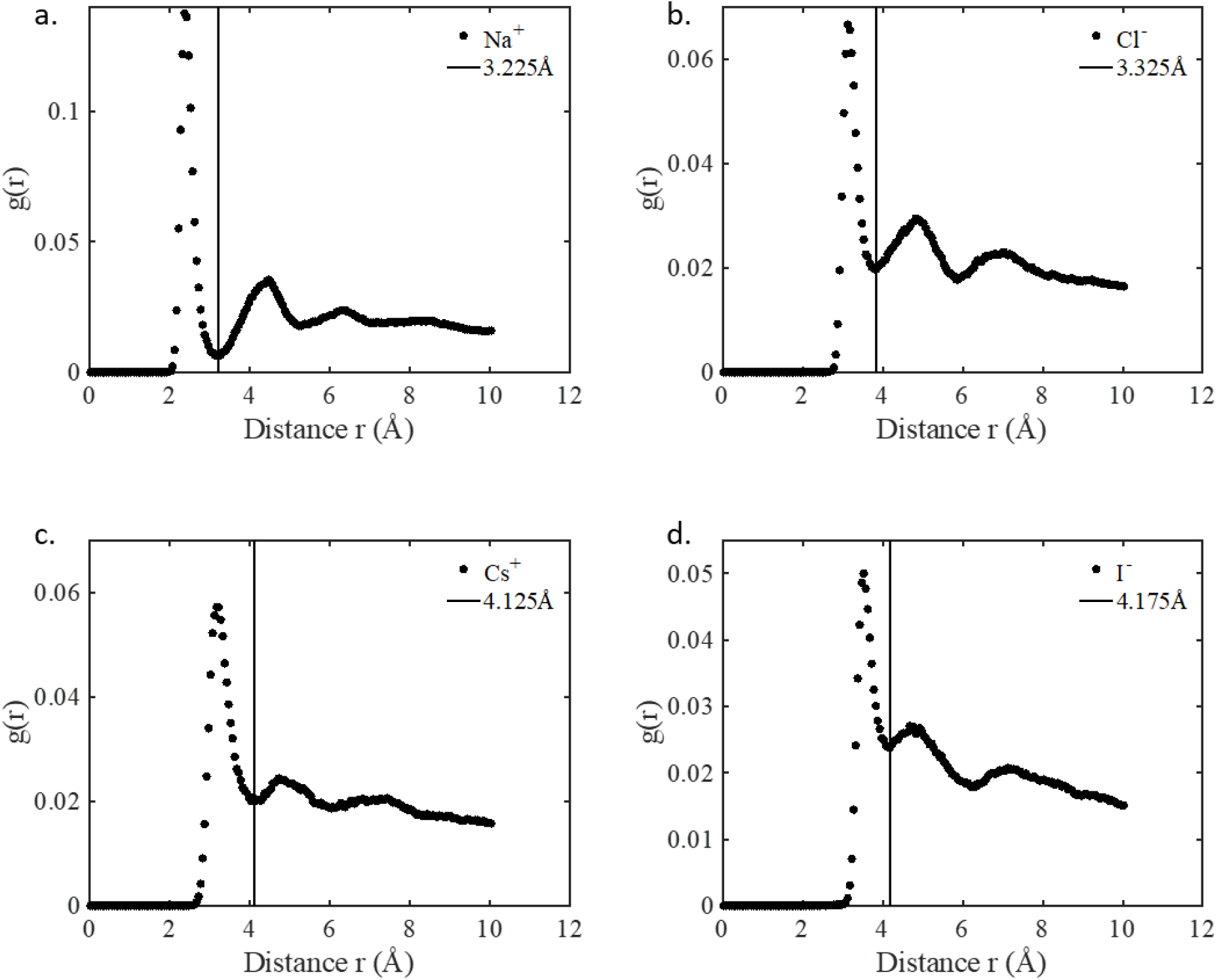
Radial pair distribution functions extracted from AIMD simulations. The first minima are used as a measure of the size of the solvation shell around different salt ions for THz data analysis (a) Na^+^, (b) Cl^-^, (c) Cs^+^, (d) I^-^. The horizontal line indicates the distance of solvation shell around each ion.

**Supplementary Figure 6.**
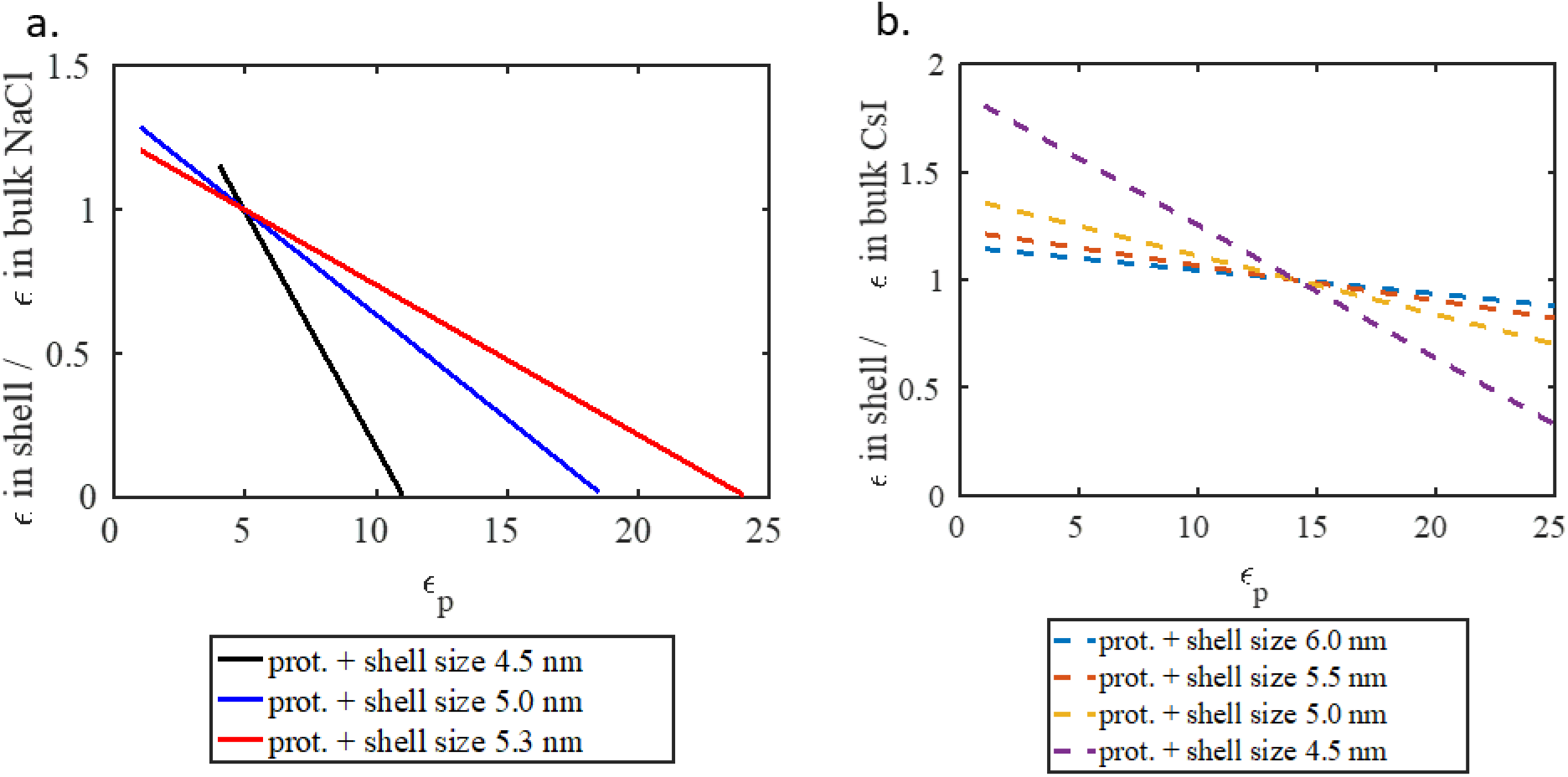
Absorption in the solvation shell excluding the protein itself is compared to bulk absorption at 1 THz. Representative ratio of the absorption in the shell compared to bulk absorption for (a) NaCl and (b) CsI for varying protein absorption (*ε*_P_) between 1-25 cm^−1^M^−1^. The solvation shell with NaCl absorbs less than the bulk above *ε*_P_ =5 cm^−1^M^−1^, while the solvation shell with CsI absorbs less than the bulk above *ε*_P_ = 15 cm^−1^M^−1^.

### Supplementary Note 2

#### Discussion of nano-ESI-MS spectra

It is likely that, at higher concentrations, even more Cs^+^ and Na^+^ ions are found to interact with aSyn, but increasing the salt concentrations in MS experiments leads to strong signal interference (Supplementary Figure 7). Relatively low intensity Na^+^ adducts can be observed in the aSyn + Cs^+^ spectra (Figure 3a and Supplementary Figure 7), the former are present in the purification buffer and are residually bound to aSyn after protein purification with a maximum of two Na^+^ bound.

**Supplementary Figure 7.**
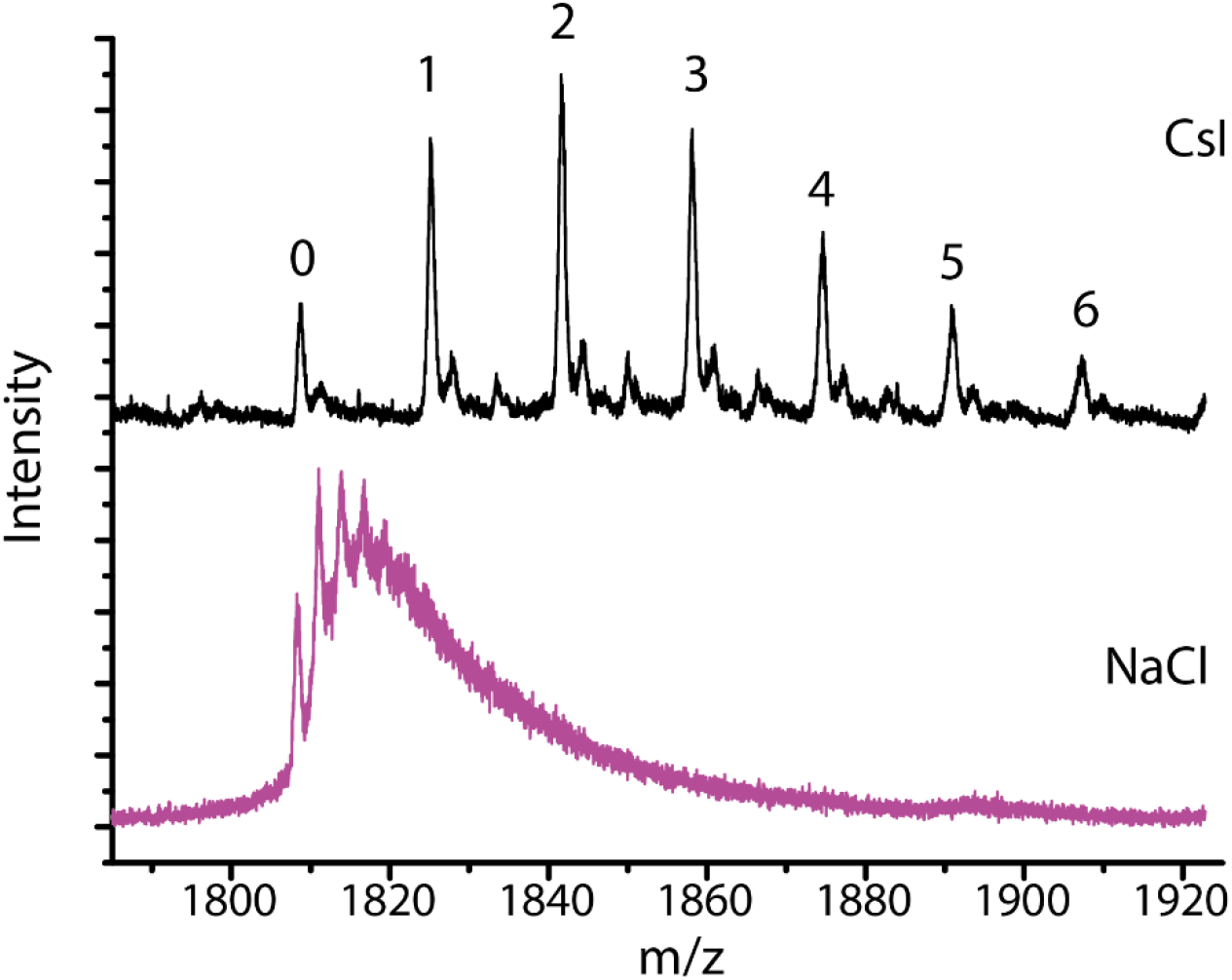
Detection of aSyn bound to NaCl and CsI at a 1:250 ratio. Nano-ESI-MS spectra in positive-ion mode for a titration of 20 μM aSyn in 5 mM NaCl (purple) and 5 mM CsI (black). There are more observed ions bound to aSyn at 5 mM compared to 1 mM (Figure 3a main text). aSyn is bound to six Cs^+^ ions at a 1:250 ratio, but for NaCl the resolution becomes distorted.

### Supplementary Note 3

#### Discussion of of nano-ESI-IM-MS spectra

The 8+ charge state was chosen to reflect one of the most physiologically relevant charge state which resembles conformations present in solution elucidated by NMR (Allison, Rivers, Christodoulou, Vendruscolo, & Dobson, 2014)^61^; this state has multiple, clearly defined conformations. Higher charge states e.g. 11+ and up correspond to more extended conformations with higher Coulombic repulsion (Supplementary Figure 8).

**Supplementary Figure 8.**
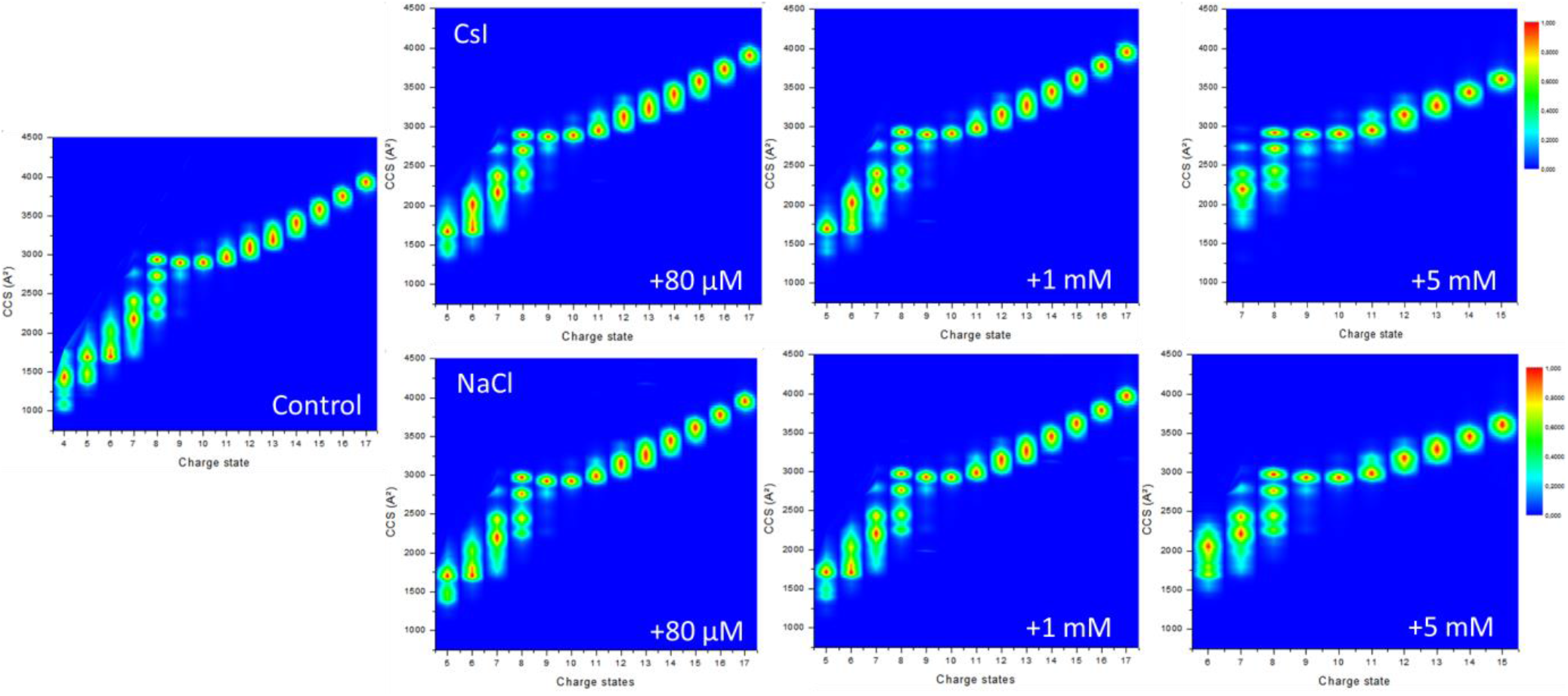
aSyn is compact at lower charge states and extended at higher charge states in the absence of and with increasing concentrations of NaCl and CsI. CCS plots of nano-ESI-IM-MS spectra of aSyn at all charge states detected. aSyn in 20 mM ammonium acetate with no salts (Control) and in the presence of 80 μM, 1 mM or 5 mM CsI and NaCl. Higher CCS values show more extended structures and extended structures are favoured at higher charge states. Shown are representative CCS plots from three injections of aSyn.

**Supplementary Figure 9.**
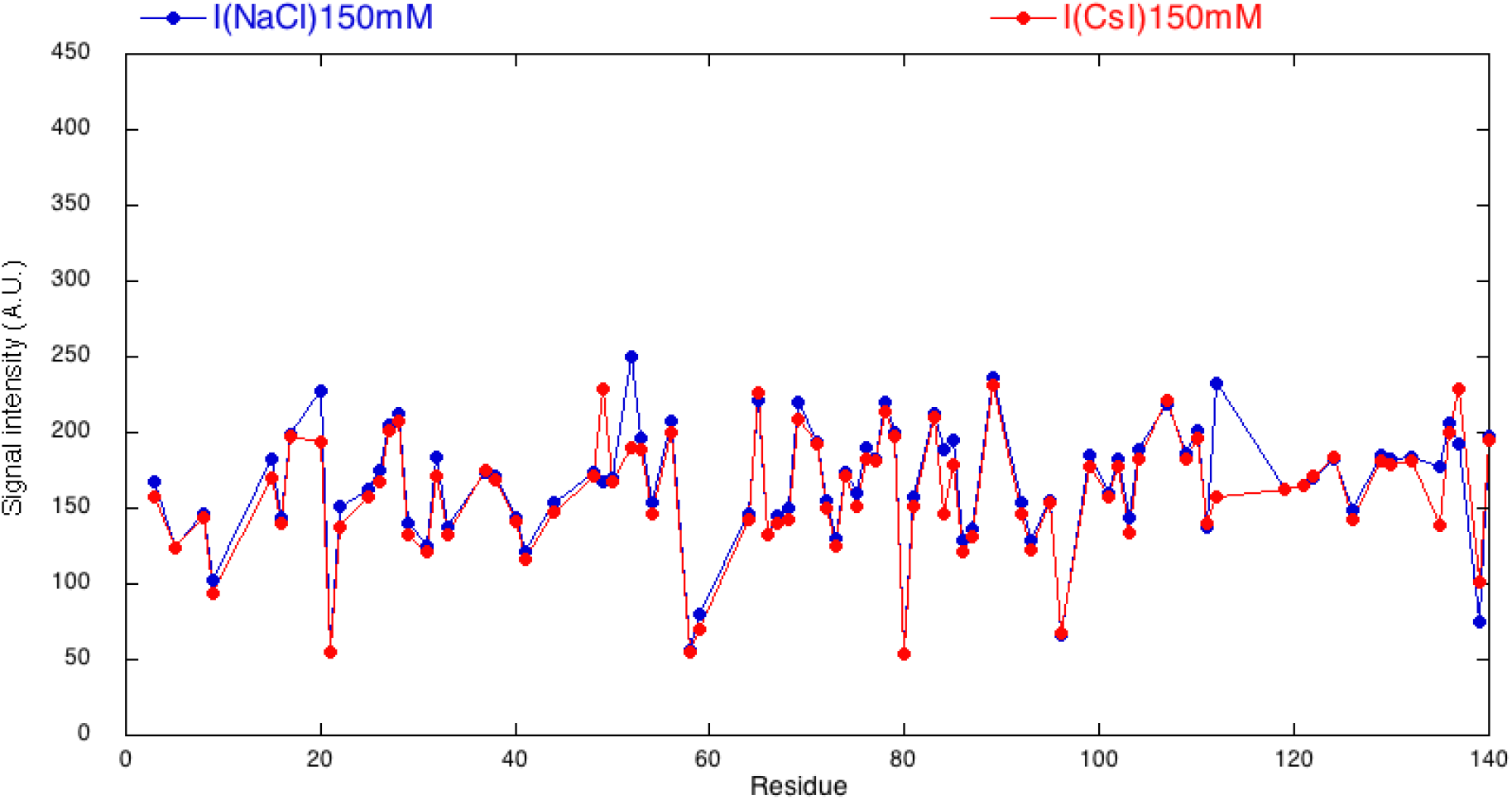
Intensity signal of ^1^H and ^15^N-labelled aSyn in 150 mM NaCl is slightly higher than in 150 mM CsI. HSQC NMR spectroscopy was used to measure the intensity of 150 μM ^1^H and ^15^N-labelled aSyn in 95% H_2_O, 5% D_2_O (vol/vol) in 150 mM CsI and NaCl. Each residue covered is represented by a spot. The signal intensity of aSyn in 150 mM NaCl is similar, but marginally high than in CsI, indicating less mobility in NaCl.

**Supplementary Figure 10.**
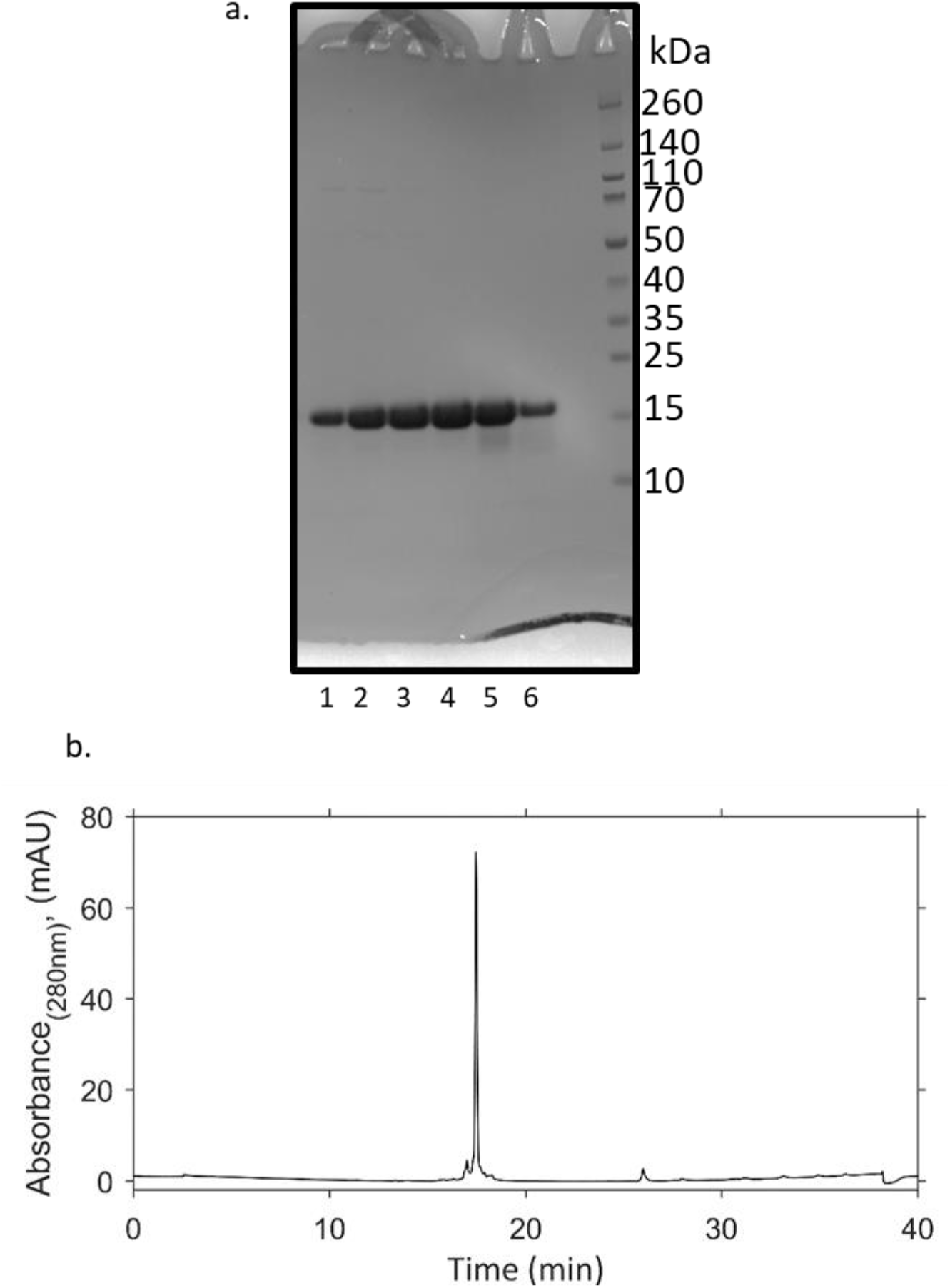
Highly pure monomeric aSyn is isolated by liquid chromatography, as shown by a representative Coomassie blue stained gel and analytical RP. (A) ^15^N-labelled aSyn after gel filtration, lanes 1-4 were used in experiments as lanes 5-6 contained degradation products. (B) 50 μL ^15^N-labelled aSyn was analysed by analytical RP-HPLC on a Discovery Bio Wide Pore C18-5 column and eluted using a gradient of 5% acetonitrile + 0.1% acetic acid to 95% acetonitrile + 0.1% acetic acid with H_2_O + 0.1% acetic acid over 40 minutes at 1 ml/min. Percentage purity of aSyn was 88.6% based on absorbance at 280 nm.

## Notes

### Competing Interest Statement

The authors have declared no competing interest.

